# Helminth-remodeled microbial indole-3-lactic acid drives AhR-dependent disease tolerance

**DOI:** 10.64898/2026.05.06.723221

**Authors:** Ruohang Sun, Yi Liu, Yang Wang, Nuo Xu, Xiaoxiao Ma, Qingbo Lv, Xiufeng Zhao, Isabelle Vallee, Pascal Boireau, Zeming Wu, Mingyuan Liu, Xiaolei Liu, Xuemin Jin

**Affiliations:** State Key Laboratory for Diagnosis and Treatment of Severe Zoonotic Infectious Diseases, Key Laboratory for Zoonosis Research of the Ministry of Education, Institute of Zoonosis, College of Veterinary Medicine, Jilin University, Changchun, 130062, China; College of Food Science and Engineering, Jilin University, Changchun, China; Würzburg Institute of Systems Immunology, Max Planck Research Group at the Julius-Maximilians University of Würzburg Würzburg, Germany; ANSES, Laboratory for Animal Health, Maisons-Alfort, France; UMR BIPAR, Anses, Laboratoire de Santé Animale, INRAE, Ecole Nationale Vétérinaire d’Alfort, Maisons-Alfort, France; School of Life Sciences, Jilin University, Changchun 130012, China

## Abstract

Helminths systemically suppress host immunity, yet whether they impose immune tolerance by rewiring host-associated microbial metabolism remains unclear. Here we show that *Trichinella spiralis* infection remodels intestinal tryptophan metabolism to generate an AhR-dependent regulatory immune state. *T. spiralis* infection enriched the commensal bacterium *Ligilactobacillus murinus*, which converted tryptophan into indole-3-lactic acid (ILA), a microbial metabolite that directly engaged the aryl hydrocarbon receptor. Antibiotic-mediated microbiota depletion abolished infection-induced ILA accumulation, AhR activation and Treg/Th17 rebalancing, whereas fecal microbiota transplantation from infected donors or supplementation with *L. murinus* or ILA restored these effects. Pharmacological blockade or genetic deletion of AhR eliminated the ability of *T. spiralis*, *L. murinus* and ILA to restrain LPS-induced cytokine-storm-like lung inflammation, establishing AhR as an essential host node in this circuit. Extending these findings to viral inflammatory disease, oral ILA improved survival and reduced pulmonary immunopathology in SARS-CoV-2-infected K18-hACE2 mice. Re-analysis of human COVID-19 metabolomic data further revealed reduced circulating ILA in severe disease. These findings define a helminth-remodeled microbial tryptophan metabolic pathway that promotes disease tolerance and identify the ILA–AhR axis as a candidate postbiotic strategy for limiting hyperinflammatory tissue injury.

## Main

Parasitic helminths are masterful manipulators of the host immune system, infecting over a billion people worldwide by actively dampening host immunity ^1^. This co-evolved tolerance strategy relies on the expansion of regulatory T cells (Tregs) and the induction of anti-inflammatory cytokines, such as IL-10 and TGF-β ^2^. The immunoregulatory capacity of helminths has been linked to the “old friends” framework, which proposes that reduced exposure to co-evolved microorganisms and metazoan parasites may contribute to the increased prevalence of immune-mediated diseases in industrialized societies. The immunoregulatory capacity of helminths has been linked to the “old friends” framework, which proposes that reduced exposure to co-evolved microorganisms and metazoan parasites may contribute to the increased prevalence of immune-mediated diseases in industrialized societies ^3–6^. Although many helminth-derived molecules act directly on host immune cells, less is known about how helminths reshape host-associated microbial metabolism to impose systemic regulatory states. Prior work has shown that the intestinal microbiota contributes to helminth-mediated suppression of allergic inflammation ^7^, yet the causal microbial taxa, metabolites, and host receptors that connect helminth exposure to immune tolerance remain incompletely defined.

The intestinal ecosystem provides a plausible route through which helminths can influence host immunity indirectly ^8,9^. ather than uniformly increasing microbial diversity, helminth infection can remodel community structure, alter the relative abundance of metabolically specialized taxa, and thereby change the pool of bioactive microbial metabolites ^9–12^. Among these pathways, microbial tryptophan metabolism is particularly relevant because gut bacteria convert dietary tryptophan into indole derivatives that function as host–microbe signaling molecules ^13,14^. Several indole metabolites activate the aryl hydrocarbon receptor (AhR), a ligand-responsive transcription factor with broad roles in barrier integrity and immune homeostasis ^15–18^. However, AhR is not a simple on–off inflammatory switch. AhR activation arbitrates the delicate balance between pro-inflammatory T helper 17 (Th17) cells and immunosuppressive Tregs. Its immunologic consequences are shaped by ligand chemistry, cellular context, tissue site, and inflammatory state. ^19,20^. This context dependence makes it essential to identify the precise microbial ligands and bacterial producers that operate during helminth infection. A key unresolved question is whether helminth-induced remodeling of the gut microbiota generates defined tryptophan-derived AhR ligands that shift the Treg/Th17 balance toward immune tolerance.

Thus, a central challenge is to move beyond correlative descriptions of helminth-associated dysbiosis and to establish a causal sequence linking a parasite-induced microbial taxon, a defined metabolite, a host receptor, and a measurable immune-protective phenotype. Here, we define a helminth–microbiota–metabolite circuit initiated by *Trichinella spiralis* infection. We show that *T. spiralis* reshapes the intestinal microbiota to enrich *Ligilactobacillus murinus*, a murine commensal capable of converting tryptophan into indole-3-lactic acid (ILA). ILA binds AhR and induces AhR-dependent transcriptional and immunologic outputs that bias the Treg/Th17 balance toward a regulatory state. Disruption of the microbiota, pharmacologic AhR blockade, or genetic AhR deficiency abrogates this regulatory program and its protection against LPS-induced cytokine-storm-like inflammation. Conversely, supplementation with either *L. murinus* or ILA recapitulates key features of helminth-induced immunoregulation. Finally, ILA improves survival and limits pulmonary inflammatory injury in SARS-CoV-2-infected K18-hACE2 mice without reducing lung viral burden, and public human metabolomic data reveal reduced circulating ILA in severe COVID-19. Together, these findings identify a helminth-remodeled microbial tryptophan metabolic pathway that promotes AhR-dependent immune tolerance and suggest that ILA-producing microbial functions, rather than helminth exposure itself, may be explored as a candidate strategy for limiting hyperinflammatory tissue injury.

## Results

### Ts infection remodeled tryptophan metabolism to modulate the Treg/Th17 balance via AhR

Tryptophan, an essential amino acid, is metabolized through three major pathways: the Kynurenine Pathway, the Serotonin Pathway, and the Indole Derivative Pathway (Fig. 1A)^15^. To investigate whether Ts infection affects host tryptophan metabolism, we performed targeted metabolomic profiling of tryptophan and its derivatives in infected mice. Principal Coordinates Analysis (PCoA) revealed a distinct separation in the metabolic profiles of fecal samples between infected and control mice (Fig. 1B and Fig. S1A), indicating a significant metabolic shift driven by the infection. The heatmap shows the relative changes in metabolite concentrations between the control group and the infection group (Fig. 1C, Fig. S1B and Table 1). We observed that Ts infection led to a significant accumulation of tryptophan in the intestine (Fig. S1C), alongside a concurrent decrease in serum tryptophan levels (Fig. S1D and Table 2). This inverse relationship suggested enhanced local tryptophan metabolism within the gut. Indeed, analysis of the three major metabolic pathways revealed that metabolites of the indole derivative pathway were significantly elevated in the feces of infected mice. Fecal levels of indole, indolelactic acid (ILA), indole-3-aldehyde (IAld), and indole-3-acetic acid (IAA) were significantly elevated. Concomitantly, serum levels of ILA were also significantly increased compared to controls (Fig. S2A-M). These data indicate that Ts infection preferentially redirects gut tryptophan metabolism toward the indole-derivative pathway, with ILA emerging as a prominent infection-associated metabolite.

**Fig 1.**
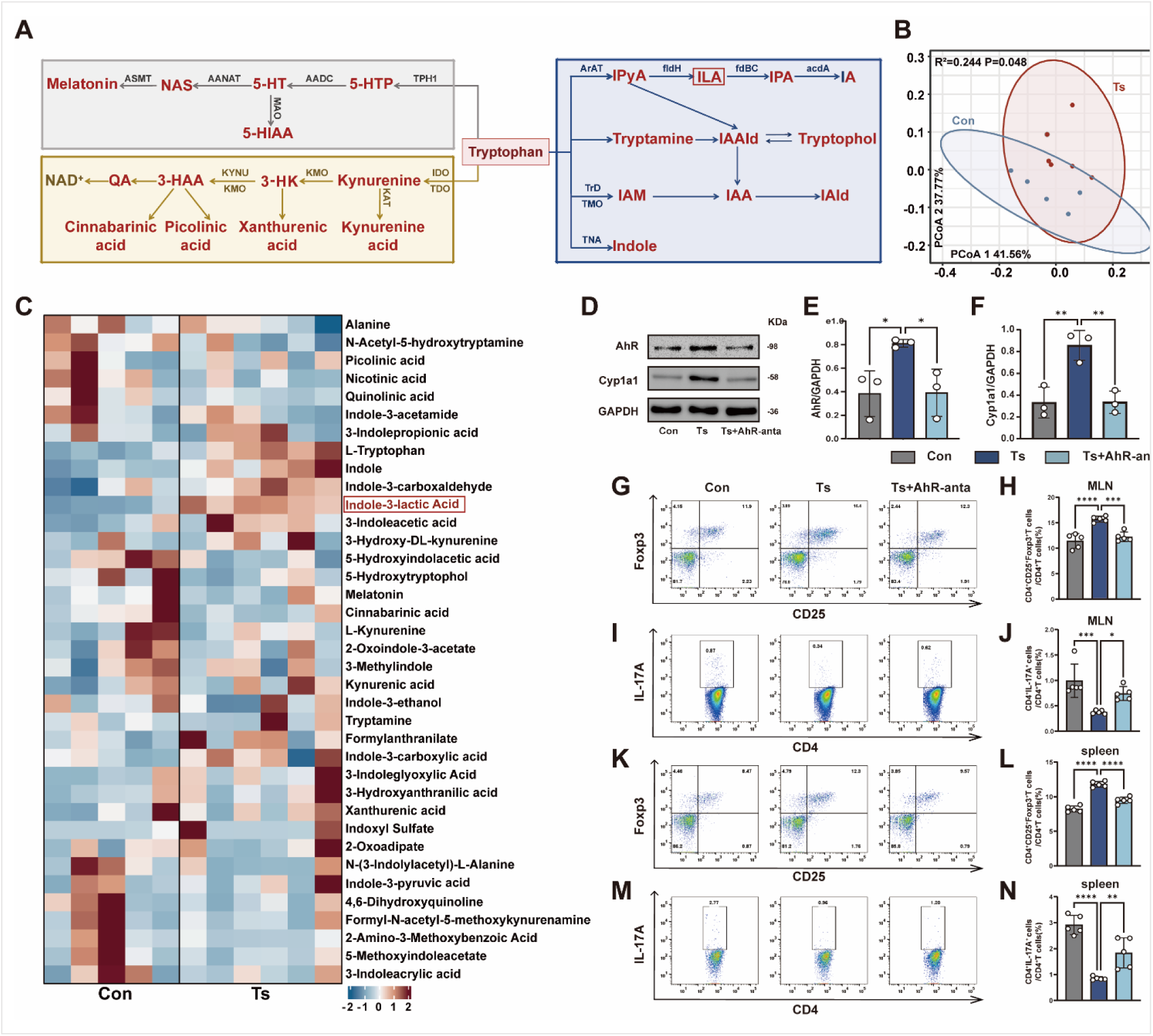
Ts infection remodels tryptophan metabolism to drive AhR-dependent modulation of the Treg/Th17 balance. **(A)** Schematic of the three major tryptophan metabolism pathways. **(B)** Principal coordinates analysis (PCoA) of fecal metabolite profiles from control and Ts-infected mice. **(C)** Heatmap of differentially abundant fecal metabolites. **(D-F)** Representative Western blot (D) and quantification (E, F) of AhR and Cyp1a1 protein level in duodenum tissues (n=3). **(G-N)** Flow cytometric analysis of Treg (CD4^+^CD25^+^Foxp3^+^) and Th17 (CD4^+^IL-17A^+^) cells in mesenteric lymph nodes (G to J) and spleen (K to N) from control, Ts-infected, and Ts-infected plus CH223191-treated mice (n=5 per group). Gating strategies are shown in Fig. S4. Data are representative of three independent experiments. Data are presented as mean ± SD. Statistical analysis was performed using one-way ANOVA with Tukey’s multiple (E, F, H, J, L and N). *p < 0.05, **p < 0.01, ***p < 0.001, ****p < 0.0001. Abbreviations: (IDO, indoleamine 2,3-dioxygenase; TDO, tryptophan 2,3-dioxygenase; KMO, kynurenine 3-monooxygenase; 3H-K, 3-hydroxykynurenine; KYNU, kynureninase; 3-HAA, 3-hydroxyanthranilic acid; QA, quinolinic acid; NAD⁺, Nicotinamide adenine dinucleotide (oxidized form); TPH1, tryptophan hydroxylase 1; AADC, aromatic L-amino acid decarboxylase; AANAT, arylalkylamine N-acetyltransferase; ASMT, Acetylserotonin O-methyltransferase; 5-HTP, 5-hydroxytryptophan; 5-HT, 5-hydroxytryptamine; MAO, monoamine oxidase; 5-HIAA, 5-hydroxyindoleacetic acid; NAS, N-acetylserotonin; ArAT, aromatic amino acid transaminase; fldH, flavin-containing L-amino acid dehydrogenase; fdBC, ferredoxin-dependent bifunctional complex; acdA, aryl-CoA dehydrogenase A; TrD, tryptophan dehydrogenase; TMO, tryptophan monooxygenase; TNA, tryptophanase; IPYA, indole-3-pyruvic acid; IPA, indole-3-propionic acid; IA, indole-3-acetic acid; IAAld, indole-3-acetaldehyde; IAM, indole-3-acetamide; IAA, indole-3-acetic acid; IAld, indole-3-aldehyde.

**Table 1.** Fecal targeted tryptophan metabolites in control and Ts-infected mice.

| Name | HMDB | Control | Ts | pvalue | FC |
| --- | --- | --- | --- | --- | --- |
| 2-Amino-3-Methoxybenzoic Acid | HMDB0060374 | 16.0800 | 5.4833 | 0.2133 | 2.9325 |
| 2-Oxoadipate | HMDB0000225 | 7168.0200 | 9160.4667 | 0.6623 | 0.7825 |
| 2-Oxoindole-3-acetate | HMDB0035514 | 50722.7000 | 46282.4500 | 0.5368 | 1.0959 |
| 3-Hydroxyanthranilic acid | HMDB0001476 | 162.3000 | 427.9167 | 0.3602 | 0.3793 |
| 3-Hydroxy-DL-kynurenine | HMDB0011631 | 316.7400 | 476.7667 | 0.4286 | 0.6644 |
| 3-Indoleacetic acid | HMDB0000197 | 2166.5000 | 3742.4500 | 0.0043 | 0.5789 |
| 3-Indoleacrylic acid | HMDB0000734 | 18.5400 | 15.8833 | 1.0000 | 1.1673 |
| 3-Indoleglyoxylic Acid | HMDB0242143 | 498.2400 | 1555.6000 | 0.0303 | 0.3203 |
| 3-Indolepropionic acid | HMDB0002302 | 1242.6600 | 1242.2500 | 1.0000 | 1.0003 |
| 3-Methylindole | HMDB0000466 | 14460.9800 | 14941.2667 | 0.9307 | 0.9679 |
| 4,6-Dihydroxyquinoline | HMDB0004077 | 80.7600 | 43.4833 | 0.5368 | 1.8573 |
| 5-Hydroxyindolacetic acid | HMDB0000763 | 465.6000 | 248.6833 | 0.0519 | 1.8723 |
| 5-Hydroxytryptophol | HMDB0001855 | 173.7800 | 113.6500 | 0.0519 | 1.5291 |
| 5-Methoxyindoleacetate | HMDB0004096 | 16.8800 | 8.6000 | 0.5830 | 1.9628 |
| Alanine | HMDB0000161 | 263526.3000 | 254010.2500 | 0.9307 | 1.0375 |
| Cinnabarinic acid | HMDB0004078 | 201.9800 | 153.9333 | 0.2468 | 1.3121 |
| Formylanthranilate | HMDB0004089 | 741.8600 | 1194.8333 | 0.3290 | 0.6209 |
| Formyl-N-acetyl-5-methoxykynurenamine | HMDB0004259 | 79.0400 | 53.8000 | 0.7922 | 1.4691 |
| Indole | HMDB0000738 | 96.7600 | 271.1333 | 0.0043 | 0.3569 |
| Indole-3-acetamide | HMDB0029739 | 8.5200 | 6.1833 | 0.6623 | 1.3779 |
| Indole-3-carboxaldehyde | HMDB0029737 | 94.8400 | 166.5500 | 0.0043 | 0.5694 |
| Indole-3-carboxylic acid | HMDB0003320 | 341.2600 | 446.7833 | 0.5368 | 0.7638 |
| Indole-3-ethanol | HMDB0003447 | 1711.4400 | 1537.6500 | 0.9307 | 1.1130 |
| Indole-3-lactic Acid | HMDB0000671 | 430.6600 | 2060.1667 | 0.0043 | 0.2090 |
| Indole-3-pyruvic acid | HMDB0060484 | 84.5800 | 96.6667 | 0.7739 | 0.8750 |
| Indoxyl Sulfate | HMDB0000682 | 1164.6200 | 14093.8333 | 0.7922 | 0.0826 |
| Kynurenic acid | HMDB0000715 | 2577.5400 | 4345.3167 | 0.2468 | 0.5932 |
| L-Kynurenine | HMDB0000684 | 249.2000 | 236.4667 | 1.0000 | 1.0538 |
| L-Tryptophan | HMDB0000929 | 61350.3600 | 115036.5000 | 0.0087 | 0.5333 |
| Melatonin | HMDB0001389 | 1.1800 | 0.9333 | 0.3580 | 1.2643 |
| N-(3-Indolylacetyl)-L-Alanine | HMDB0304378 | 44.1400 | 27.6000 | 0.2923 | 1.5993 |
| N-Acetyl-5-hydroxytryptamine | HMDB0001238 | 120.0200 | 59.9500 | 0.0823 | 2.0020 |
| Nicotinic acid | HMDB0001488 | 45004.7400 | 25831.2833 | 0.2468 | 1.7423 |
| Picolinic acid | HMDB0002243 | 68551.6200 | 59280.8500 | 0.9307 | 1.1564 |
| Quinolinic acid | HMDB0000232 | 2778.7400 | 1134.9167 | 0.2468 | 2.4484 |
| Tryptamine | HMDB0000303 | 1183.1000 | 1853.3667 | 1.0000 | 0.6384 |
| Xanthurenic acid | HMDB0000881 | 1844.4800 | 3137.7333 | 0.4286 | 0.5878 |

**Table 2.** Serum targeted tryptophan metabolites in control and Ts-infected mice.

| Name | HMDB | Control | Ts | pvalue | FC |
| --- | --- | --- | --- | --- | --- |
| Indoxylsulfate | HMDB0000682 | 5844.5779 | 9484.0288 | 0.3939 | 0.6163 |
| 5-Hydroxyindoleacetic acid | HMDB0000763 | 119.1303 | 149.1278 | 0.1797 | 0.7988 |
| N-acetylserotonin | HMDB0001238 | 5.4704 | 4.4062 | 0.5887 | 1.2415 |
| 3-Indoleacrylic acid | HMDB0000734 | 4.5310 | 18.7826 | 0.0043 | 0.2412 |
| Xanthurenic acid | HMDB0000881 | 22.6163 | 8.8625 | 0.0022 | 2.5519 |
| Indole-3-lactic acid | HMDB0000671 | 154.3958 | 284.0603 | 0.0260 | 0.5435 |
| Indolylpropionic acid | HMDB0002302 | 402.6000 | 841.1525 | 0.0087 | 0.4786 |
| L-tryptophan | HMDB0000929 | 25176.1464 | 11151.8267 | 0.0022 | 2.2576 |
| β-Indole-3-acetic acid | HMDB0000197 | 86.9914 | 149.5702 | 0.2403 | 0.5816 |
| Hydroxytryptophan | HMDB0000472 | 60.1390 | 46.8825 | 0.1320 | 1.2828 |
| Serotonin | HMDB0000259 | 2074.4299 | 1810.3482 | 0.8182 | 1.1459 |
| 2-Ketoadipic acid | HMDB0000225 | 430.7010 | 184.7024 | 0.0087 | 2.3319 |
| 3-Hydroxykynurenine | HMDB0000732 | 13.5827 | 9.7941 | 0.0931 | 1.3868 |
| L-kynurenine | HMDB0000684 | 151.6479 | 261.9623 | 0.0411 | 0.5789 |
| Nicotinic acid | HMDB0001488 | 31.9785 | 26.5733 | 0.1320 | 1.2034 |
| 2-Aminobenzoic acid | HMDB0001123 | 6.4422 | 10.4118 | 0.0649 | 0.6187 |
| Indole-3-acetamide | HMDB0029739 | 8.1693 | 1.3977 | 0.0022 | 5.8448 |
| Indole-3-carbaldehyde | HMDB0029737 | 20.8223 | 5.2479 | 0.0022 | 3.9678 |
| Picolinic acid | HMDB0002243 | 10.5261 | 6.9685 | 0.0649 | 1.5105 |
| N-Formylanthranilic acid | HMDB0004089 | 3.1584 | 1.8379 | 0.2403 | 1.7184 |
| N-formylkynurenine | HMDB0001200 | 6.8620 | 1.6273 | 0.0048 | 4.2169 |

Tryptophan metabolites can act as endogenous ligands to activate the AhR. As a key physiological regulator of immune homeostasis, AhR activation modulates the balance between Treg and Th17 cells.^21^. Disruption of the Treg/Th17 cells balance is strongly associated with autoimmune diseases and systemic inflammation. We explored whether Ts can activate the AhR signaling pathway. AhR protein and mRNA expression levels were significantly elevated in intestine of mice infected with Ts. Cytochrome P450 Family 1 Subfamily A Member 1 (Cyp1a1) serves as both a key downstream effector of AhR and a classical biomarker for AhR activation. Further analysis of Cyp1a1 expression revealed that Ts infection significantly upregulated both protein and mRNA levels of Cyp1a1. (Fig. 1D-F and Fig. S3A, B). We next investigated if Ts infection modulates host immunity via the AhR signaling pathway. Compared to controls, Ts infection significantly increased the mRNA expression of the Treg-associated transcription factor Foxp3 and the immunosuppressive cytokine TGF-β, while downregulating the Th17-associated transcription factor RORγt and its effector cytokine IL-17A (Fig. S3, C-F). Pharmacological AhR blockade reversed these transcriptional and cellular changes, supporting an AhR-dependent contribution to the Ts-induced shift in the Treg/Th17 balance. Furthermore, flow cytometric analysis of lymphocyte subsets in the mesenteric lymph nodes (MLNs) and spleen revealed that the proportion of CD4⁺CD25^+^Foxp3⁺ Treg cells was significantly increased, while the proportion of CD4⁺IL-17A⁺ Th17 cells was markedly decreased in the infected group (Fig. 1G-N, and Fig. S4). AhR antagonist treatment reversed these changes, confirming the central role of the AhR pathway in this immunomodulation.

### Ts-induced alteration in tryptophan metabolism was microbiota dependent

The gut microbiota has been confirmed as a critical modulator of tryptophan metabolism^22^. To verify the mediating role of the gut microbiota, we established an antibiotic-treated (ABX) mouse model with depleted commensal bacteria followed by Ts infection (Fig. 2A). Metagenomic analysis confirmed that ABX treatment reduced α-diversity and markedly altered β-diversity, consistent with effective microbiota depletion (Fig. S5A-C). In contrast to microbiota-intact mice, Ts infection failed to increase fecal tryptophan-derived indoles in ABX-treated mice; fecal tryptophan and ILA levels remained low and were not significantly different between ABX and ABX+Ts groups (Fig. 2B, C, fig. S5D, E, and Table 3). Meanwhile, Ts infection failed to activate the AhR signaling pathway in microbiota depletion, as no infection-induced upregulation of AhR or Cyp1a1 was observed in the intestinal tissue (Fig. 2D-F, and fig. S5F, G). Crucially, microbiota depletion abolished the Ts-induced shift in the Treg/Th17 balance (Fig. 2G-J), highlighting the essential role of the gut microbiota. To test if the helminth-remodeled microbiota was sufficient to mediate these effects, we performed a fecal microbiota transplantation (FMT) experiment (Fig. 2K). Feces from uninfected control donors or donors infected with Ts for 14 days were transplanted into ABX recipient mice. We confirmed that there was no worm burden in the intestine of Ts-infected mice at 14 days post infection, as we have documented in previous study ^23,24^. Microscopic examination confirmed that no live adult worms or larvae were detectable in the fecal inocula used for FMT (data not shown). With this experiment, the influence of helminth can be ruled out through transplanting the feces from mice at this stage of infection. The results showed that no significant differences in the expression levels of AhR and Cyp1a1 were observed in the FMT-con group compared with the control group, while the expression levels of AhR and Cyp1a1 were significantly increased in the FMT-Ts group compared with both the control and FMT-con groups (Fig. 2L-N, and Fig. S5H, I). Moreover, mRNA expression of Foxp3 and TGF-β was markedly upregulated, while RORγt and IL-17A expression was significantly downregulated compared to the control group (Fig. 2O-R). Collectively, these findings indicate that the gut microbiota serves as a crucial intermediary in Ts-mediated regulation of host tryptophan metabolism and the Treg/Th17 immune balance. Because the infection-remodeled microbiota generated higher levels of infection-associated indole metabolites than the baseline microbiota (Fig. 1 and Fig. S2), we next sought to identify the bacterial taxa linked to this metabolic shift.

**Fig 2.**
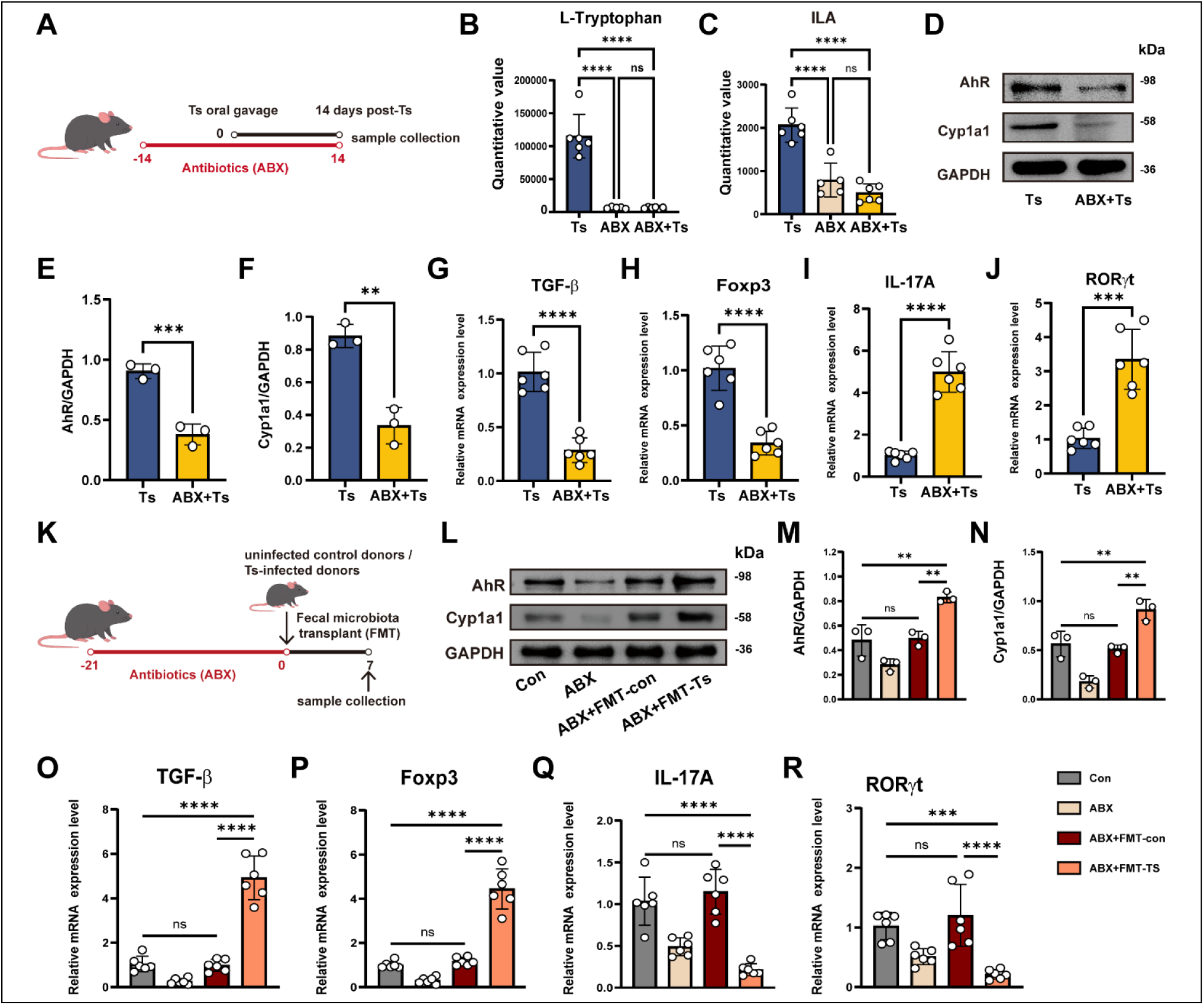
Ts-driven immunomodulation is indirect and mediated by the gut microbiota. **(A-J)** Gut microbiota depletion abolishes Ts-induced AhR activation and immunomodulation. (A) Schematic of the antibiotic (ABX) treatment experiment (ampicillin 0.5 g/L, neomycin 0.5 g/L, metronidazole 0.5 g/L, vancomycin 0.25 g/L). (B, C) Fecal levels of L-Tryptophan and ILA (n=6). (D-F) Western blot analysis of AhR and Cyp1a1 protein expression in the duodenum (n=3). (G-J) qPCR analysis of Treg/Th17-associated gene expression in the duodenum (n=6). **(K-R)** Fecal microbiota transplantation (FMT) from Ts-infected donors (250 TsML) recapitulates AhR activation and immunomodulation. (K) Schematic of the FMT experiment. (L-N) Western blot analysis of AhR and Cyp1a1 protein expression in the duodenum (n=3). (O-R) qPCR analysis of Treg/Th17-associated gene expression in the duodenum (n=6). Data are representative of three independent experiments. Data are presented as mean ± SD and were analyzed by Student’s t-test (E-J) or one-way ANOVA with Tukey’s test (B, C and M-R). **p < 0.01, ***p < 0.001, ****p < 0.0001.

**Table 3.** Fecal targeted tryptophan metabolites in ABX and ABX+Ts mice.

| Name | HMDB | ABX | ABX+Ts | pvalue | FC |
| --- | --- | --- | --- | --- | --- |
| Alanine | HMDB0000161 | 69117.5459 | 64911.3635 | 0.6623 | 1.0648 |
| L-Tryptophan | HMDB0000929 | 6527.3996 | 6623.7212 | 1.0000 | 0.9855 |
| L-Kynurenine | HMDB0000684 | 280.3456 | 419.7025 | 0.0173 | 0.6680 |
| 3-Indolepropionic acid | HMDB0002302 | 43.3563 | 50.5600 | 0.5368 | 0.8575 |
| 3-Hydroxy-DL-kynurenine | HMDB0011631 | 392.7803 | 444.7782 | 0.1775 | 0.8831 |
| Nicotinic acid | HMDB0001488 | 6611.5928 | 4304.0672 | 0.0087 | 1.5361 |
| Kynurenic acid | HMDB0000715 | 331.0456 | 925.2977 | 0.0043 | 0.3578 |
| 5-Hydroxyindolacetic acid | HMDB0000763 | 256.1505 | 323.5625 | 0.7922 | 0.7917 |
| 3-Indoleacetic acid | HMDB0000197 | 164.7936 | 91.5851 | 0.1775 | 1.7993 |
| Xanthurenic acid | HMDB0000881 | 1614.7126 | 1254.3929 | 0.1255 | 1.2872 |
| N-Formylkynurenine | HMDB0001200 | 504.8974 | 780.1546 | 0.4286 | 0.6472 |
| Picolinic acid | HMDB0002243 | 848.2722 | 937.6252 | 0.3290 | 0.9047 |

| **Name** | **HMDB** | **ABX** | **ABX+Ts** | **pvalue** | **FC** |
| --- | --- | --- | --- | --- | --- |
| N-Acetyl-5-hydroxytryptamine | HMDB0001238 | 1.3032 | 1.6304 | 0.6623 | 0.7993 |
| Indole-3-acetamide | HMDB0029739 | 0.5732 | 0.5743 | 1.0000 | 0.9981 |
| Indole | HMDB0000738 | 144.6844 | 109.2351 | 0.2468 | 1.3245 |
| Indole-3-ethanol | HMDB0003447 | 187.6695 | 291.1229 | 0.3290 | 0.6446 |
| Nicotinamide | HMDB0001406 | 85.7041 | 99.3099 | 0.3290 | 0.8630 |
| 5-Methoxytryptamine | HMDB0004095 | 2.1415 | 1.4734 | 0.1699 | 1.4535 |
| 3-Methylindole | HMDB0000466 | 732.0122 | 466.9759 | 0.6623 | 1.5676 |
| 6-Hydroxymelatonin | HMDB0004081 | 1.3079 | 1.7711 | 0.4102 | 0.7385 |
| Melatonin | HMDB0001389 | 1.2139 | 0.7837 | 0.1699 | 1.5489 |
| Tryptamine | HMDB0000303 | 5.6572 | 6.9891 | 0.2468 | 0.8094 |
| Serotonin | HMDB0000259 | 1305.1474 | 1736.3428 | 0.3290 | 0.7517 |
| 5-Hydroxytryptophol | HMDB0001855 | 165.1548 | 263.3214 | 0.0303 | 0.6272 |
| N-Methyltryptamine | HMDB0004370 | 0.1869 | 0.1960 | 0.9271 | 0.9535 |
| Indole-3-carboxylic acid | HMDB0003320 | 104.0676 | 78.6972 | 0.1255 | 1.3224 |
| Indole-3-lactic Acid | HMDB0000671 | 789.1117 | 493.1209 | 0.1775 | 1.6002 |
| 3-Hydroxyanthranilic acid | HMDB0001476 | 556.4710 | 500.8588 | 0.9271 | 1.1110 |
| Quinolinic acid | HMDB0000232 | 520.1405 | 340.2686 | 0.1255 | 1.5286 |
| 2-Oxoadipate | HMDB0000225 | 11766.3206 | 14093.0560 | 0.1775 | 0.8349 |
| Formylanthranilate | HMDB0004089 | 83.7359 | 102.9958 | 0.3290 | 0.8130 |
| 3-Indoleglyoxylic Acid | HMDB0242143 | 482.9229 | 399.7825 | 0.2468 | 1.2080 |
| Formyl-N-acetyl-5-methoxykynurenamine | HMDB0004259 | 22.5244 | 7.2069 | 0.5368 | 3.1254 |
| Indole-3-carboxaldehyde | HMDB0029737 | 63.1553 | 42.3842 | 0.0519 | 1.4901 |
| Indole-3-pyruvic acid | HMDB0060484 | 98.8899 | 87.7484 | 0.4286 | 1.1270 |
| 2-Oxoindole-3-acetate | HMDB0035514 | 7294.9272 | 3949.2349 | 0.5368 | 1.8472 |
| Indoxyl Sulfate | HMDB0000682 | 2713.3128 | 217.4851 | 0.3290 | 12.4759 |
| 4,6-Dihydroxyquinoline | HMDB0004077 | 112.7724 | 62.8504 | 0.0173 | 1.7943 |

### Ts infection enriched for *L. murinus*, which produced the direct AhR ligand ILA

To identify the microbial taxa responsible for the observed metabolic changes, we performed metagenomic sequencing of fecal samples. The results showed that, compared to the control group, there were no significant differences in richness indices (Chao1 and observed species) or diversity indices (Simpson and Shannon) in the gut microbiota of Ts-infected mice (Fig. S6A). However, Venn diagram analysis revealed a clear distinction in gut microbial composition between the different groups (Fig. S6B). Principal coordinate analysis (PCoA) based on taxonomic abundance profiles revealed a significant separation between control and Ts-infected mice (Fig. 3A). At the phylum level, the infected group showed an increase in the abundance of *Bacillota* and *Verrucomicrobia*. At the family level, gut microbiota remodeling in the infected group was characterized by elevated abundances of *Akkermansiaceae*, *Bacteroidaceae*, and *Lactobacillaceae* (Fig. S6C). At the genus level, we observed an increase in the abundance of *Ligilactobacillus*, *Lactobacillus*, and *Akkermansia* in the infected group (Fig. 3B). We performed Linear Discriminant Analysis Effect Size (LEfSe) with an LDA score threshold > 3. Notably, *L. murinus* showed the largest LDA effect size among taxa enriched in Ts-infected mice, nominating it as a candidate microbial contributor to the altered tryptophan-metabolic profile (Fig. 3C, D). Spearman correlation analysis revealed a strong positive association between the abundance of *L. murinus* and fecal ILA levels in Ts-infected mice (Fig. 3E), indicating that Ts-induced alterations in the gut microbiota contribute to indole metabolism. In addition, Wilck et al. demonstrated that germ-free mice mono-colonized with *L. murinus* exhibited increased fecal tryptophan metabolites (ILA, IAld, IAA) versus control germ-free mice^25^. To confirm that *L. murinus* directly produces ILA from tryptophan, we analyzed the supernatant of *L. murinus* cultures. Our results confirmed that *L. murinus* is capable of metabolizing tryptophan into ILA (Fig. 3F-G, and Table 4). Taken together, our findings suggest that Ts promotes tryptophan metabolism and elevates ILA levels by increasing the abundance of *L. murinus* in the host gut.

**Fig 3.**
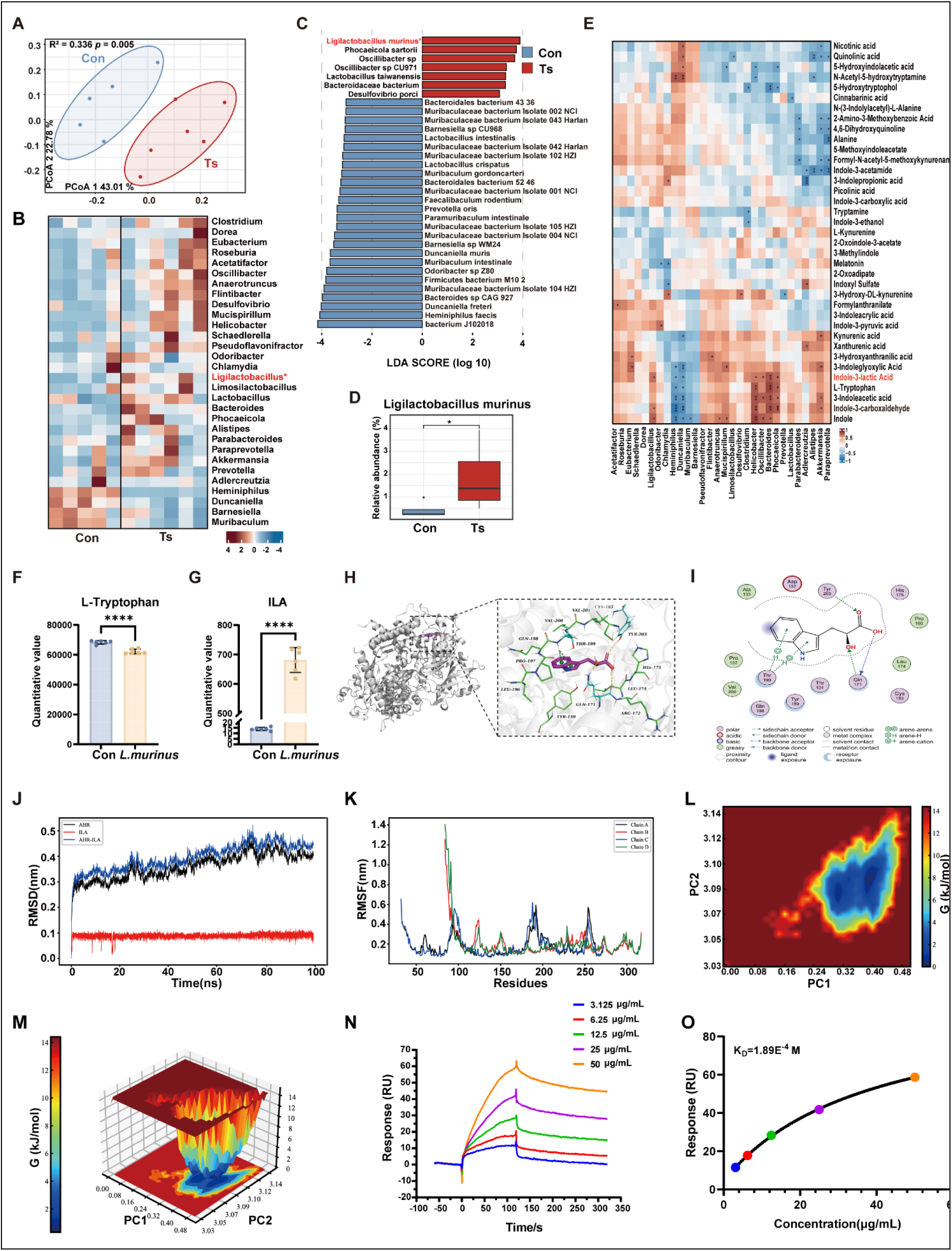
Ts infection enriches for *L. murinus*, which produces the direct AhR ligand ILA. **(A)** PCoA plot showing distinct gut microbial composition in Ts-infected mice. **(B)** Heatmap of relative bacterial abundance at the genus level. **(C)** LEfSe analysis identifying enriched bacterial taxa (LDA score > 3, Kruskal-Wallis test) **(D)** Relative abundance of *L. murinus* in control and Ts-infected mice. **(E)** Spearman correlation matrix linking bacterial taxa with tryptophan metabolites. **(F, G)** Validation of ILA production from tryptophan by *L. murinus* in vitro (n=6). **(H-M)** Molecular docking and dynamics simulations demonstrating stable binding of ILA to the AhR ligand-binding pocket. (J, K) Root-mean-square deviation (RMSD) and fluctuation (RMSF) plots indicating system stability. **(N, O)** Surface Plasmon Resonance (SPR) analysis quantifying the direct binding affinity between ILA and recombinant AhR. The sensorogram shows binding responses at graded concentrations. Note: Concentrations were prepared in μg/mL (3.125, 6.25, 12.5, 25, 50). Steady-state affinity fitting yields a calculated K_D_ of 1.89E^−4^M. Data are presented as mean ± SD. Statistical analysis for metagenomic data (C) was performed using the Kruskal-Wallis test. Comparisons in (D, F, and G) were analyzed using an unpaired Student’s t-test. *p < 0.05, ****p < 0.0001.

**Table 4.**
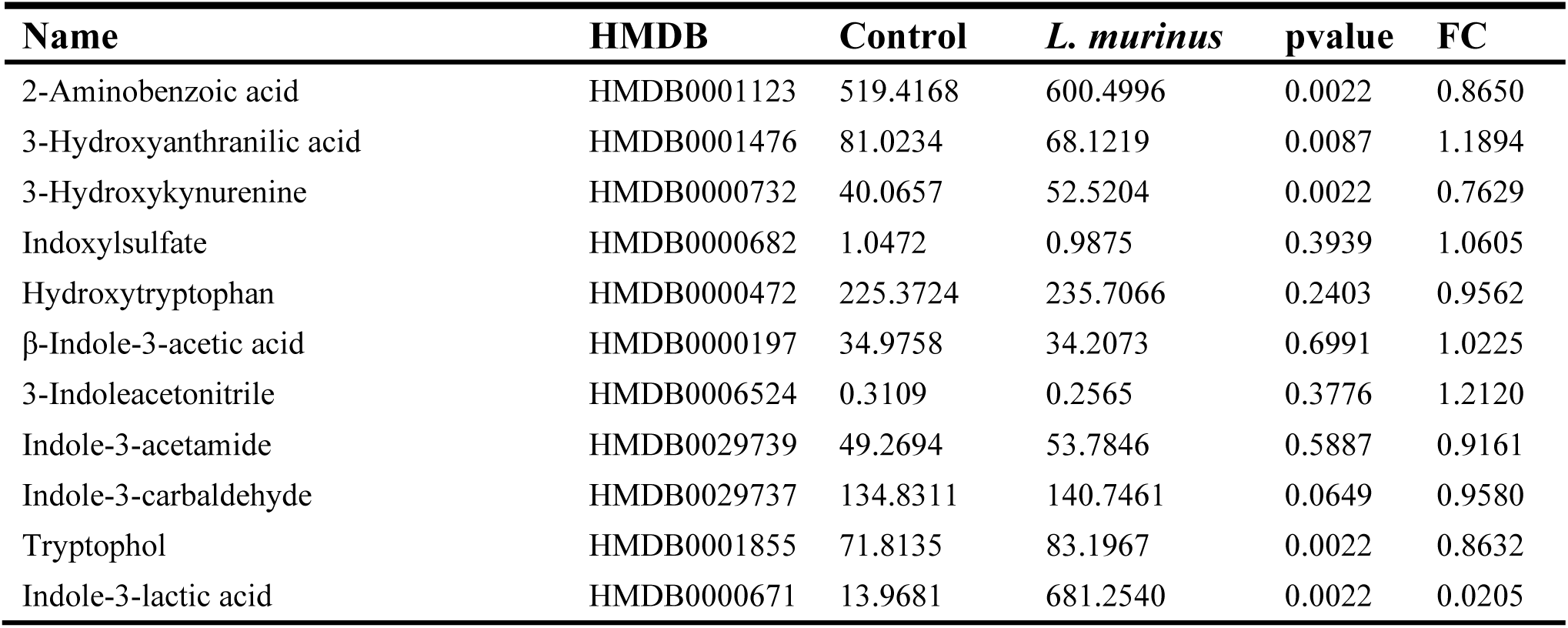
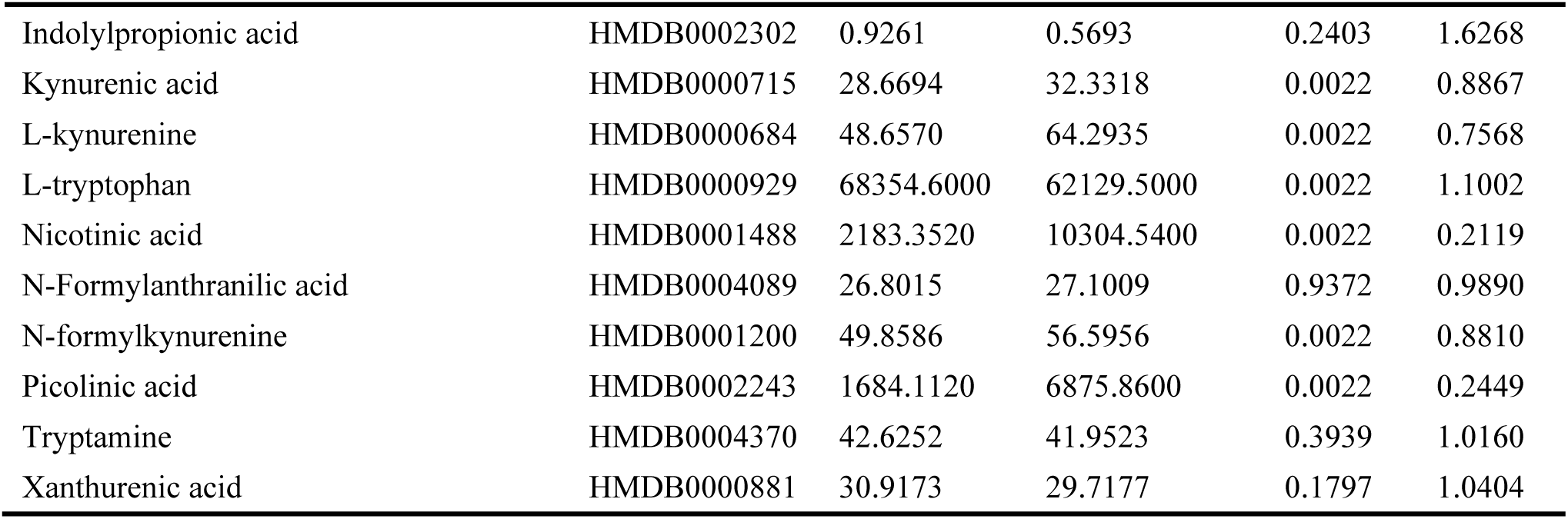
Tryptophan metabolite levels in the culture supernatant of *L. murinus*.

To elucidate the molecular basis of the interaction between ILA and the host aryl hydrocarbon receptor (AhR), we first employed molecular docking to simulate the binding of ILA within the Per-Arnt-Sim (PAS)-B domain, the primary ligand-binding pocket of AhR^19^. The computational model predicted that ILA fits precisely into the hydrophobic pocket, forming a stable hydrogen-bonding network with key residues Thr199, Gln171, and Tyr203 (Fig. 3H-I). The calculated binding energy of −6.7 kcal·mol⁻¹ suggested a high-fidelity interaction, supporting ILA as a direct, physiologically relevant but relatively low-affinity microbial ligand for AhR. To further evaluate the dynamic stability of this interaction under physiological conditions, we performed 100 ns molecular dynamics (MD) simulations. The root-mean-square deviation (RMSD) and fluctuation (RMSF) profiles reached convergence quickly and remained stable throughout the trajectory, indicating that ILA binding induces a compact and robust conformation of the AhR molecule (Fig. 3J-K). Importantly, we constructed the Free Energy Landscape (FEL) based on the MD trajectories to explore the thermodynamic properties of the complex. The FEL analysis identified a distinct, well-defined energy minimum (deep blue basin), confirming that ILA binding stabilizes AhR in a thermodynamically favored state, which is a hallmark of effective receptor activation (Fig. 3L-M). We then performed Surface Plasmon Resonance (SPR) analysis using recombinant human AhR to provide definitive biophysical evidence of direct binding. ILA exhibited a dose-dependent binding response with a calculated equilibrium dissociation constant (K_D_) of 1.89E^−4^M (Fig. 3N-O). Although this affinity is substantially weaker than that of high-affinity xenobiotic AhR ligands, it supports direct engagement of AhR by ILA within a concentration range that may be relevant for locally abundant microbial metabolites.

### *L. murinus* and ILA supplementation recapitulated AhR activation and Treg/Th17 modulation

Having identified *L. murinus* and its metabolite ILA as key mediators, we next tested whether their administration alone could recapitulate the immunomodulatory effects of Ts infection in antibiotic treated mice. We examined whether L. murinus supplementation affected intestinal adult worm burden during Ts infection. Mice were orally administered *L. murinus* continuously for 14 days after infection, and the intestinal adult worm burden was assessed at 14 dpi. Compared with the control group, *L. murinus* supplementation significantly increased the intestinal adult worm burden (Fig. 4A). After depleting the gut microbiota in mice using antibiotic treatment, the experimental groups were supplemented with *L. murinus*. After continuous intervention for 7 days, relevant indicators in intestinal tissues were assessed. Compared to vehicle controls, supplementation with *L. murinus* significantly increased both mRNA and protein expression of AhR and Cyp1a1 in the intestine (Fig. 4B-D and Fig. S7 A, B). Additionally, mRNA expression of Foxp3 and TGF-β was significantly upregulated in the intervention groups, while expression of RORγt and IL-17A was markedly suppressed compared to the control group (Fig. S7C). Flow cytometric analysis further demonstrated that *L. murinus* supplementation significantly increased the proportion of CD4⁺CD25⁺Foxp3⁺ Treg cells while reducing the proportion of CD4⁺IL-17A⁺ Th17 cells in the mesenteric lymph nodes (MLNs) (Fig. 4E–H). Similar results were also observed in the flow cytometric analysis of the spleen, where *L. murinus* markedly promoted Treg cell expansion and suppressed Th17 cell differentiation (Fig. 4I–L). These findings indicate that *L. murinus* activates the AhR signaling pathway and shifts the Treg/Th17 balance toward immune tolerance. Similarly, following antibiotic treatment, mice were supplemented with ILA. Intestinal adult worm burden analysis showed that, compared with the Ts-infected group, ILA supplementation also significantly increased the small intestinal adult worm burden at 14 dpi (Fig. 4M). Compared with vehicle controls, ILA significantly enhanced both the mRNA and protein expression levels of AhR and Cyp1a1 in intestinal tissues (Fig. 4N–P and Fig. S7D, E). qPCR analysis further confirmed that ILA significantly upregulated the mRNA expression of Foxp3 and TGF-β while suppressing the expression of RORγt and IL-17A (Fig. S7F). Flow cytometric analysis showed that ILA significantly increased the proportion of CD4⁺CD25⁺Foxp3⁺ Treg cells and decreased the proportion of CD4⁺IL-17A⁺ Th17 cells in both the MLNs and spleen (Fig. 4 Q–X). Overall, gut microbiota-derived indole metabolites activate the AhR pathway, increasing Treg frequencies while reducing Th17 frequencies. Previous studies have shown that Treg-mediated immune tolerance facilitates parasite persistence by suppressing host anti-parasitic immunity^26^, which may explain the increased intestinal adult worm burden following *L. murinus* and ILA supplementation. In vitro stimulation of intestinal organoids with *L. murinus* or ILA significantly upregulated mRNA expression levels of AhR and Cyp1a1 in a dose-dependent manner, as determined by qPCR analysis (Fig. S7G-L). In intestinal organoids, *L. murinus* or ILA increased AhR and Cyp1a1 expression, and this induction was attenuated by AhR antagonist treatment, indicating that the *L. murinus*–ILA axis can activate AhR signaling in the intestinal epithelial compartment (Fig. 4Y).

**Fig 4.**
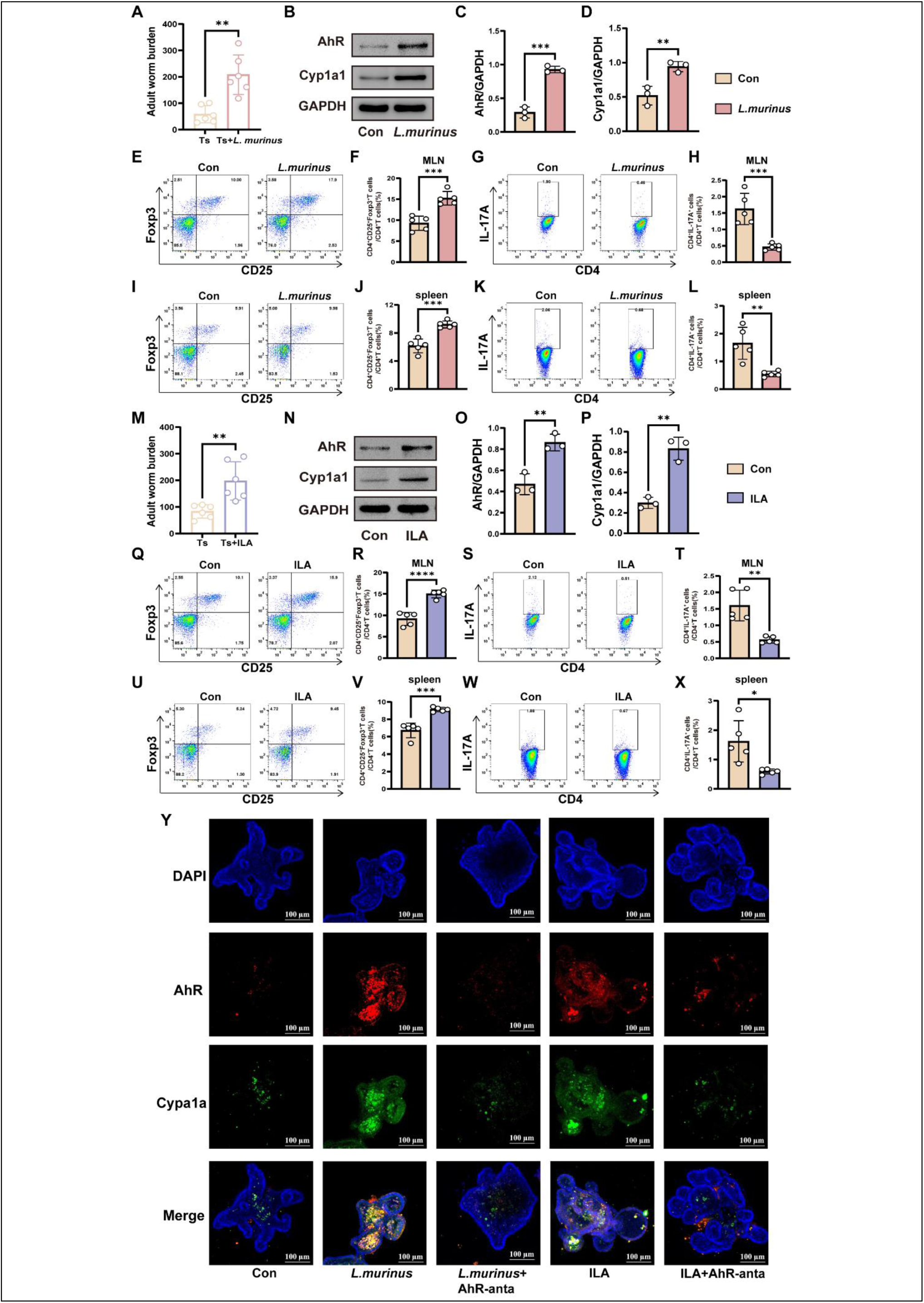
Supplementation with *L. murinus* or ILA recapitulates AhR activation and immunomodulation. **(A)** Mice were orally infected with 500 TsML, followed by oral administration of vehicle control or *L. murinus* (1 × 10^9 CFU/mL) from 1 to 14 dpi, and at 14 dpi, adult worms were recovered from each group to assess the burden of *T. spiralis* (n=6). **(B-D)** In vivo effects in antibiotic-treated mice. Western blot analysis and quantification of AhR and Cyp1a1 protein levels in the duodenum after supplementation with vehicle, *L. murinus* (1×10^9^ CFU/mL) for 7days (n=3). **(E-L)** Flow cytometric analysis of Treg (CD4^+^CD25^+^Foxp3^+^) and Th17 (CD4^+^IL-17A^+^) cells in mesenteric lymph nodes (E, F, G and H) and spleen (I, J, K and L) after supplementation with vehicle, *L. murinus* (1×10^9^ CFU/mL) for 7days after 21 days of antibiotic pretreatment (n=5). **(M)** Mice were orally infected with 500 TsML, followed by oral administration of vehicle control or ILA (20 mg/kg) from 1 to 14 dpi, and at 14 dpi, adult worms were recovered from each group to assess the burden of *T. spiralis* (n=6). **(N-P)** In vivo effects in antibiotic-treated mice. (N-P) Western blot analysis and quantification of AhR and Cyp1a1 protein levels in the duodenum after supplementation with vehicle, ILA (20 mg/kg) for 7days (n=3). **(Q-X)** Flow cytometric analysis of Treg (CD4^+^CD25^+^Foxp3^+^) and Th17 (CD4^+^IL-17A^+^) cells in mesenteric lymph nodes (Q, R, S and T) and spleen (U, V, W and X) after supplementation with vehicle, ILA (20 mg/kg) for 7days after 21 days of antibiotic pretreatment (n=5). **(Y)** Representative immunofluorescence images confirming increased AhR and Cyp1a1 protein in organoids treated with *L. murinus* (1×10^6^ CFU/mL) or ILA (2 mM) for 48 h. Data are representative of three independent experiments. Data are presented as mean ± SD and were analyzed by Student’s t-test (A, C, D, F, H, J, L, M, O, P, R, T, V, and X). *p < 0.05, **p < 0.01, ***p < 0.001, ****p < 0.0001.

### Ts-*L. murinus*-ILA Axis conferred AhR-dependent protection against LPS-induced cytokine storm

To determine whether the Ts-*L. murinus*-ILA axis confers functional protection against hyperinflammation, we used a murine model of LPS-induced cytokine storm. Ts infection markedly attenuated this inflammatory syndrome, as reflected by reduced LPS-induced lung pathology and inflammatory cell infiltration (Fig. 5A, B). Moreover, it lowered serum levels of the pro-inflammatory cytokines tumor necrosis factor-α (TNF-α) and interferon-γ (IFN-γ), increased the Treg-associated anti-inflammatory cytokines IL-10 and TGF-β, and decreased the Th17-associated cytokine IL-17A (Fig. 5C-G). This protection was entirely AhR-dependent, as it was abolished by pretreatment with the AhR antagonist CH223191. Previous characterization has confirmed that CH223191 lacks agonist activity and does not induce downstream signaling or toxicity in the absence of exogenous ligands ^27^, indicating that the observed loss of protection is due to specific signaling blockade. In line with our hypothesis, supplementation with either *L. murinus* or ILA recapitulated the protective effects of Ts infection, significantly reducing lung histopathology scores and inflammatory infiltration in the LPS-induced cytokine storm model (Fig. 5H, I). Serum levels of TNF-α, IFN-γ, and IL-17A were decreased, while IL-10 and TGF-β levels were elevated, closely mirroring the immunoregulatory profile induced by Ts infection (Fig. 5J-N). Similarly, pretreatment with CH223191 also abolished the protective effects of *L. murinus* and ILA. Genetic ablation of AhR provided definitive confirmation of its central role. In AhR-knockout (AhR-KO) mice, the protective effects of Ts infection, *L. murinus*, and ILA against LPS-induced lung pathology and cytokine storm were completely abolished. (Fig. 5O-U).

**Fig 5.**
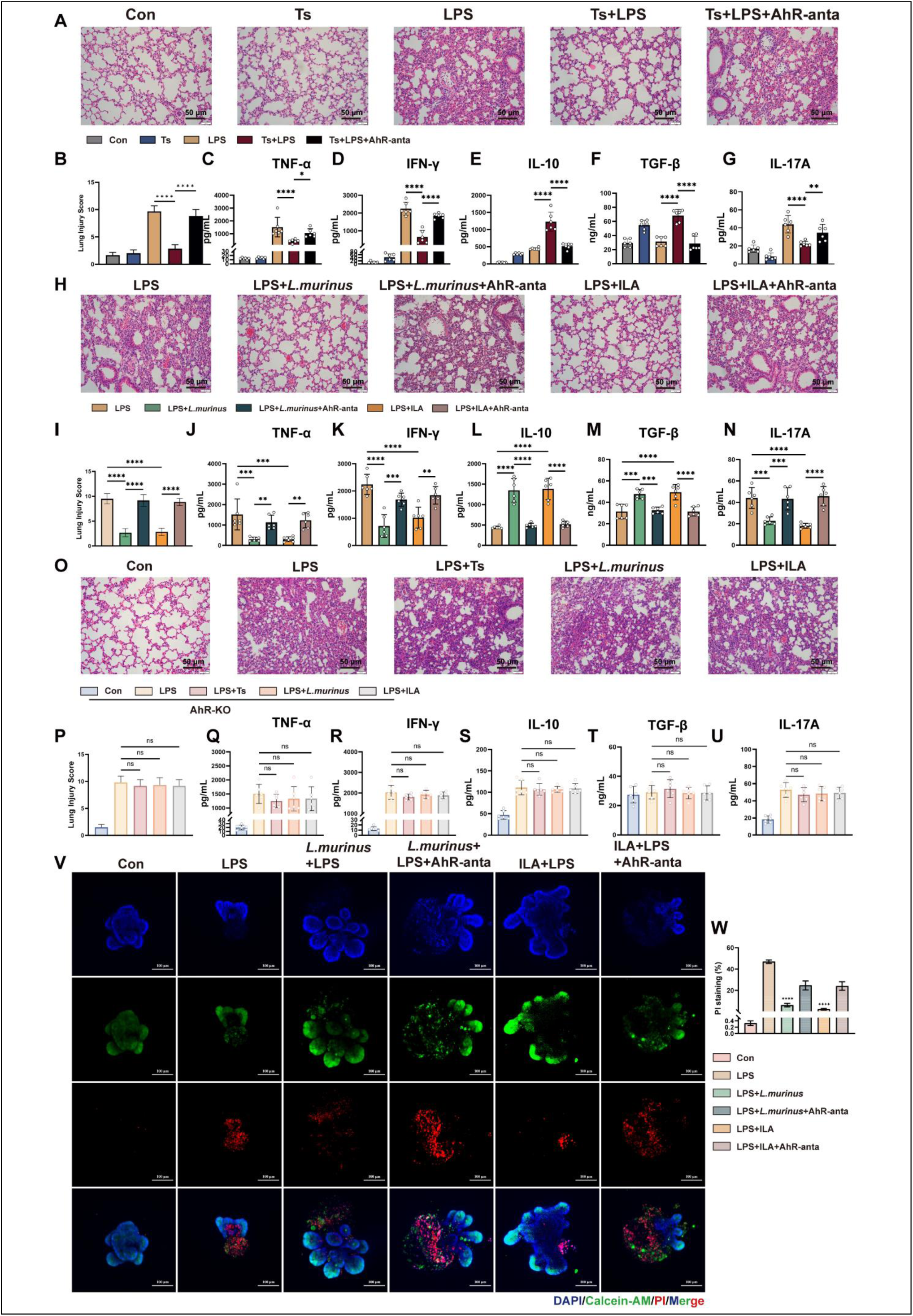
The Ts-*L. murinus*-ILA axis protects against LPS-induced cytokine storm through AhR. **(A-G)** Ts infection protects wild-type (WT) mice from LPS (8 mg/kg) -induced cytokine storm, an effect reversed by the AhR antagonist CH223191 (10 mg/kg). (A) Representative H&E staining of lung sections. (B) Lung histopathological scores. (C-G) Serum cytokine levels. **(H-N)** Supplementation with *L. murinus* (1×10^9^ CFU/mL) or ILA (20 mg/kg) mimics the protective effects in WT mice. (H) Representative lung H&E staining. (I) Lung scores. (J-N) Serum cytokine levels. **(O-U)** The protective effects of Ts infection, *L. murinus*, or ILA are completely abolished in AhR-knockout (AhR-KO) mice. (O) Representative lung H&E staining. (P) Lung scores. (Q-U) Serum cytokine levels. **(V, W)** *L. murinus* and ILA protect intestinal organoids from LPS-induced damage in vitro. (V) Data are representative of three independent experiments. Representative images of organoids with PI staining indicating cell death. (W) Quantification of damaged organoids. n=6 for in vivo experiments and n=3 for in vitro experiments. Scale bars, 100 μm. Data are presented as mean ± SD. Statistical analysis was performed using one-way ANOVA with Tukey’s multiple comparison test (B, C, E, F, H, I and K). ns, no statistical significance, *p < 0.05, ***p < 0.001, ****p < 0.0001.

In vitro, LPS treatment increased PI-positive cell death and reduced organoid viability, whereas *L. murinus* or ILA improved viability in an AhR-dependent manner (Fig. 5V, W). QPCR and ELISA results further demonstrated decreased expression of pro-inflammatory cytokines TNF-α and IFN-γ (Fig. S8A-D). Notably, this protective effect was abolished by CH223191 treatment. Collectively, these findings indicate that Ts exerts anti-inflammatory and protective effects by activating the AhR pathway through the *L. murinus*-ILA axis, thereby modulating the Treg/Th17 cells balance.

### ILA improves disease outcome after lethal SARS-CoV-2 challenge

Cytokine storm represents a significant cause of morbidity and mortality in patients with COVID-19 ^28^. We further evaluated whether ILA could alter disease outcome after SARS-CoV-2 infection (Fig. 6A). ILA treatment markedly improved disease outcomes in SARS-CoV-2-infected K18-hACE2 mice. While SARS-CoV-2 infection caused rapid mortality, with all mice succumbing by 9 dpi, oral administration of ILA substantially increased survival, with approximately 60% of mice surviving to 14 dpi (Fig. 6B). Consistently, infected mice showed progressive body-weight loss, whereas ILA-treated mice exhibited significantly attenuated weight loss and partial recovery during the later phase of infection (Fig. 6C). At 6 dpi, pulmonary viral load was comparable between SARS-CoV-2-infected mice with or without ILA treatment, indicating that ILA did not significantly reduce lung viral burden (Fig. 6D). However, ILA markedly reshaped the pulmonary cytokine response. SARS-CoV-2 infection increased lung TNF-α, IFN-γ, and IL-17A levels, whereas ILA treatment significantly reduced these pro-inflammatory cytokines (Fig. 6E, F and I). In contrast, ILA strongly enhanced IL-10 production and moderately increased TGF-β levels in infected lungs (Fig. 6G-H). The improvements were observed in the histopathology in the lungs of the mice treated with ILA (Fig. 6J and K). Together, these results indicate that ILA protects SARS-CoV-2-infected K18-hACE2 mice without directly suppressing viral replication, but instead by limiting excessive pulmonary inflammation and promoting a regulatory cytokine profile.

**Fig 6.**
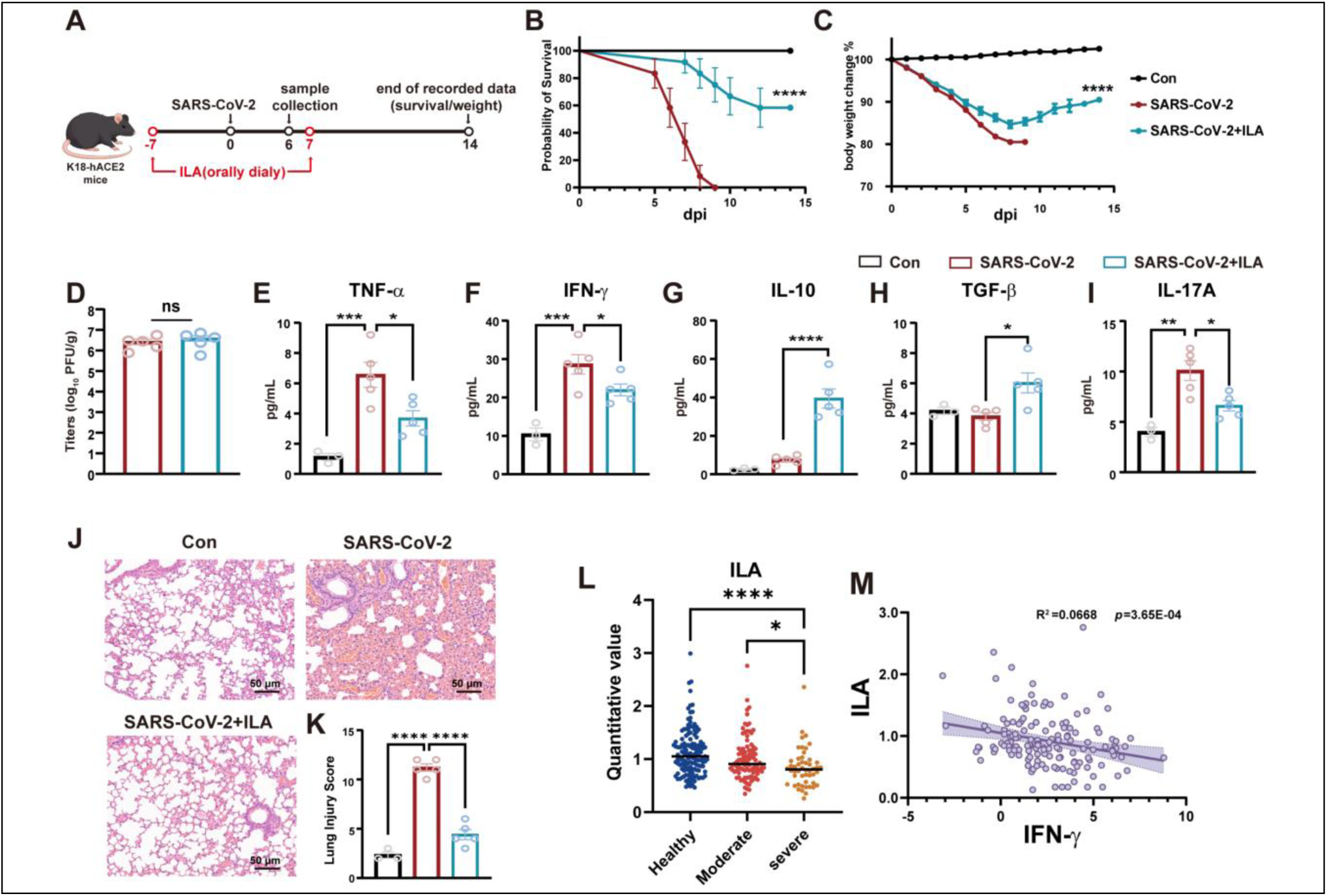
ILA protects against SARS-CoV-2 lethality and cytokine storm in mice and humans. **(A)** Experimental design to determine whether ILA (20 mg/kg) protected against SARS-CoV-2 infection in 5- to 6-week-old K18-hACE2 mice. **(B)** Survival of ILA-treated K18-hACE2 mice after SARS-CoV-2 infection (n =12). **(C)** Percent changes of body weights of ILA-treated K18-hACE2 mice after SARS-CoV-2 infection (n=12). **(D)** Lung viral load of mice after SARS-CoV-2 infection (n=5). **(E-I)** Serum cytokine levels (n=3-5). **(J)** Representative H&E staining of lung sections (n=3-5). **(K)** Lung histopathological scores (n=3-5). **(L)** Serum level of ILA is progressively reduced with increasing disease severity. **(M)** Scatterplots demonstrating a significant negative correlation between serum level of ILA and the pro-inflammatory cytokine IFN-γ. Data were re-analyzed from a published cohort (Su et al., 2020) ^29^ comprising healthy controls (n=258), patients with moderate COVID-19 (n=102), and patients with severe COVID-19 (n=52), the latter representing the subgroup most consistent with a cytokine storm phenotype. Statistical analysis was performed using the survival curve comparison (log-rank [Mantel-Cox] test) (B), Student’s t-test (D) and one-way ANOVA with Tukey’s multiple comparison test (C, E, F, G, H, I, K and L). ns, no statistical significance, *p < 0.05, **p < 0.01, ***p < 0.001, ****p < 0.0001. Linear regression trend lines are shown with 95% confidence intervals indicated by shading (M).

To bridge our murine findings to human disease, we asked whether a similar deficiency in the ILA-AhR axis is associated with cytokine storm-related hyperinflammation in humans. In a previously published study by Su et al ^29^. proteomic and metabolomic profiling was performed on serum samples from 102 patients with moderate COVID-19, 52 patients with severe COVID-19, and 258 healthy controls. We re-analyzed the metabolomic dataset from this cohort, focusing on moderate-to-severe COVID-19, particularly the severe subgroup characterized by a cytokine storm phenotype. Serum ILA levels were reduced in patients with COVID-19, with the lowest levels observed in the severe group (Fig 6L). Consistent with the protection against LPS- or SARS-CoV-2-induced cytokine storm observed in mice, circulating ILA level was inversely correlated with the pro-inflammatory cytokine IFN-γ in these patients (Fig 6M). These findings suggest that depletion of protective indole metabolites may be linked to the cytokine storm state in severe COVID-19.

## Discussion

Helminths have long been viewed as direct immunological manipulators that deploy parasite-derived molecules to restrain host inflammatory responses. Our study expands this view by identifying a microbial-metabolic route through which *Trichinella spiralis* imposes systemic immune tolerance. We show that *T. spiralis* infection remodels intestinal tryptophan metabolism, enriches the ILA-producing commensal *L. murinus*, activates AhR signaling, and shifts the Treg/Th17 balance toward a regulatory state (Fig 7). This pathway was not merely correlative: antibiotic-mediated microbiota depletion abolished the metabolic and immunological effects of infection, fecal transfer from infected donors reproduced key features of the phenotype, and supplementation with either *L. murinus* or ILA was sufficient to recapitulate AhR activation and immune regulation. These findings position helminths not only as sources of immunomodulatory molecules, but also as ecological engineers of host-associated microbial metabolism. In this model, a metazoan parasite rewires a bacterial metabolic circuit to generate a host-active ligand that restrains inflammatory pathology.

**Fig 7.**
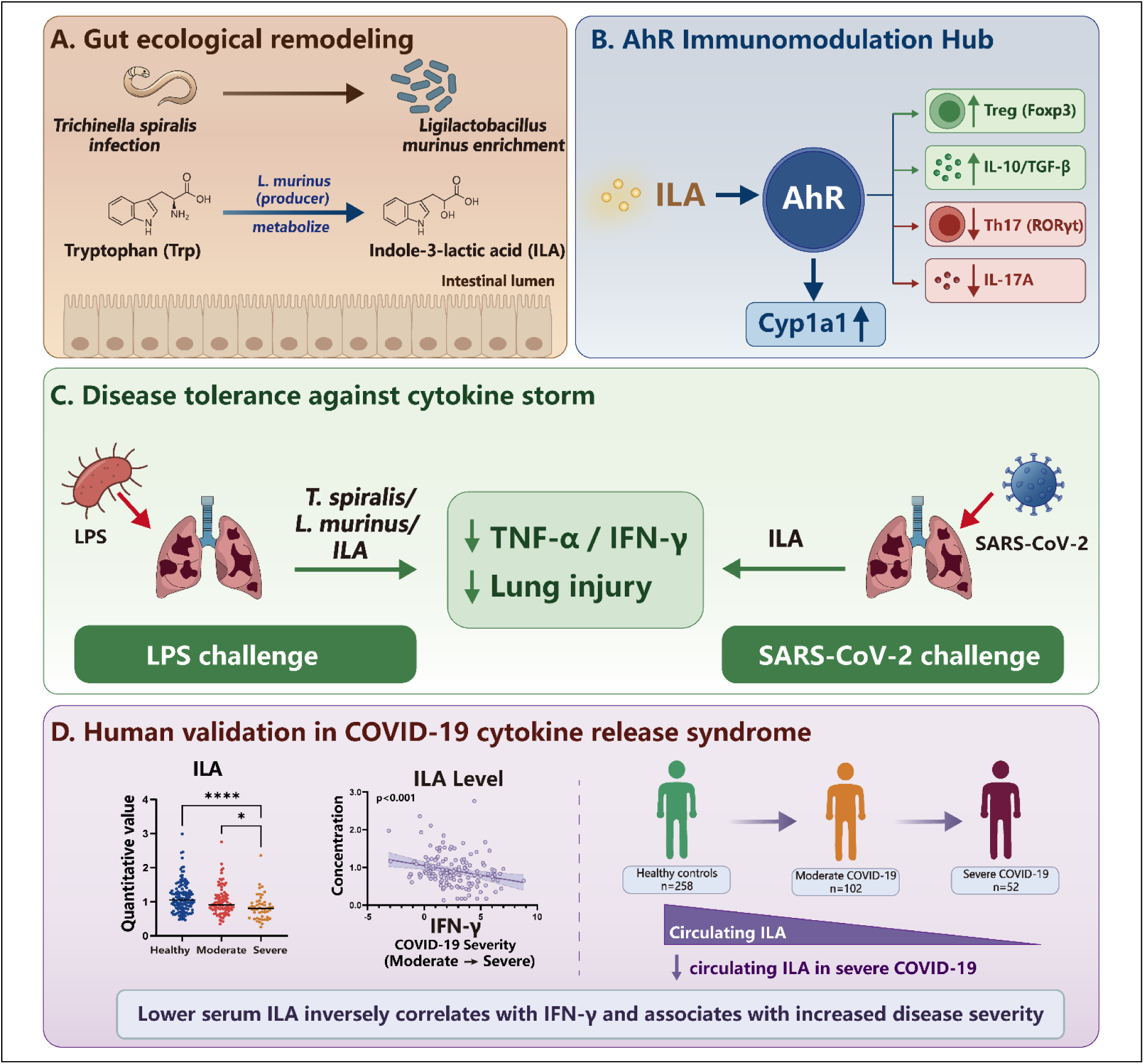
Proposed model of the helminth-remodeled microbial ILA-AhR axis in disease tolerance. **(A)** *Trichinella spiralis* infection remodels the gut ecological niche and enriches the ILA-producing commensal *Ligilactobacillus murinus*. *L. murinus* metabolizes tryptophan into indole-3-lactic acid (ILA) in the intestinal lumen. **(B)** Microbial ILA engages the aryl hydrocarbon receptor (AhR), induces the canonical AhR target Cyp1a1, and establishes an immunoregulatory hub characterized by increased Treg/Foxp3 and IL-10/TGF-β responses, together with reduced Th17/RORγt and IL-17A responses. **(C)** In experimental cytokine-storm models, *T. spiralis* infection, *L. murinus*, or ILA limits LPS-induced pulmonary inflammation, while ILA also attenuates SARS-CoV-2-induced lung injury, reducing inflammatory cytokines such as TNF-α and IFN-γ. **(D)** Re-analysis of human COVID-19 metabolomic data shows that circulating ILA levels are reduced in severe disease and inversely associated with IFN-γ, linking impaired ILA availability to COVID-19 cytokine release syndrome and disease severity.

A central advance of this work is the identification of a defined causal chain linking a parasite-induced bacterial taxon, a microbial metabolite, a host receptor, and a disease-relevant immune phenotype. Previous studies have shown that helminths can alter the intestinal microbiota and that the microbiota contributes to helminth-mediated suppression of inflammatory disease ^7^. However, the specific bacterial functions and host receptors connecting helminth-induced dysbiosis to immune tolerance have remained incompletely resolved. Here, *L. murinus* emerged as a major infection-enriched taxon associated with increased fecal ILA. Culture-based metabolite profiling further confirmed that *L. murinus* can convert tryptophan into ILA, while bacterial or metabolite supplementation reproduced the major immunological features of infection. These findings are in line with previous reports identifying ILA as a bioactive microbial metabolite produced by commensal bacteria, including *Lactobacillus* species ^30,31^. This interpretation is also compatible with recent evidence that *L. murinus*-derived tryptophan metabolites help sustain fetomaternal tolerance ^32^, suggesting that this metabolic module may represent a broader host-microbe mechanism for maintaining immune homeostasis across distinct physiological contexts. These findings suggest that the immunoregulatory outcome of helminth infection is not dictated solely by the presence of the parasite, but by the metabolic state of the intestinal ecosystem that the parasite helps create ^15,33^.

Among the tryptophan-derived metabolites altered by *T. spiralis* infection, ILA was the most extensively validated functional mediator in this study. Microbial tryptophan catabolites are increasingly recognized as interkingdom signaling molecules that shape barrier and immune homeostasis ^15^. Our data extend this concept by placing ILA within a helminth-remodeled immune circuit. ILA directly engaged AhR in biophysical and computational analyses, Notably, the direct interaction between ILA and AhR appears to be of relatively low affinity compared with classical high-affinity xenobiotic AhR ligands. This feature may better fit the biology of microbiota-derived metabolites, which often act as locally abundant, context-dependent physiological ligands that tune barrier and immune homeostasis without necessitating sustained high-amplitude receptor activation. In this framework, the protective effect of ILA is consistent with promotion of disease tolerance ^34^. ILA induced canonical AhR-responsive programs in intestinal tissue and organoids, and promoted a regulatory immune profile characterized by increased Foxp3, TGF-β, IL-10 and Treg frequencies, together with reduced RORγt, IL-17A and Th17 responses. Importantly, pharmacologic AhR blockade and genetic AhR deficiency abolished the protective effects of *T. spiralis*, *L. murinus* and ILA, establishing AhR as an essential host node in this pathway. These results also reinforce a broader principle of AhR biology: AhR is not a simple pro- or anti-inflammatory switch, but a ligand-, cell-and context-dependent interpreter of environmental and microbial signals ^30,33^. Structurally distinct AhR ligands can elicit divergent transcriptional outcomes depending on tissue site, inflammatory state and cellular target. In the present setting, ILA appears to favor an immunoregulatory output rather than an overtly inflammatory program.

Functionally, the *T. spiralis*–*L. murinus*–ILA–AhR axis limited cytokine-storm-like pathology in two inflammatory settings. In the LPS model, helminth infection, *L. murinus* and ILA each reduced lung injury and pro-inflammatory cytokine production in an AhR-dependent manner. In SARS-CoV-2-infected K18-hACE2 mice, ILA improved survival and reduced pulmonary inflammatory injury without significantly lowering lung viral burden. This distinction is important. The protective phenotype is better interpreted as enhanced disease tolerance, or limitation of immune-mediated tissue damage, rather than improved pathogen resistance. Such a tolerance-like mechanism may be particularly relevant to viral or sterile inflammatory conditions in which mortality is driven not only by pathogen burden, but also by excessive host inflammatory responses. ILA treatment protected SARS-CoV-2-infected mice from lethal inflammatory disease, and re-analysis of a human multi-omics cohort revealed lower circulating ILA in severe COVID-19, with an inverse association between ILA and IFN-γ. These observations are consistent with reports linking disturbed microbial tryptophan metabolism to acute and post-acute SARS-CoV-2 inflammatory outcomes ^29,35^. Future studies should test whether ILA abundance, ILA-producing microbial capacity or AhR-responsive transcriptional signatures prospectively predict inflammatory severity and treatment response in human disease.

The evolutionary implications of this pathway are notable. ILA and *L. murinus* increased intestinal adult worm burden, indicating that the regulatory state induced by this axis may facilitate parasite persistence. Rather than representing a purely host-protective pathway, the *L. murinus*–ILA–AhR circuit may therefore reflect a negotiated tolerance state: the parasite benefits from restrained anti-helminth immunity, whereas the host benefits from reduced inflammatory damage. This trade-off is conceptually important for therapeutic development. Enhancing ILA-AhR signaling may be beneficial in cytokine-driven tissue injury, but could be detrimental in settings where strong type 2 immunity, Th17 immunity or pathogen clearance is required. Therapeutic exploitation of this pathway will therefore require careful attention to disease context, timing, dose, tissue compartment and infection status.

Several limitations must be acknowledged to guide future research and eventual translation to human health. First, a key limitation is the species-specificity, as *L. murinus* is not a major constituent of the human gut microbiota. This underscores the need to shift from a “species-centric” to a “function-centric” approach, as the therapeutic principle lies in the metabolic function—ILA production—rather than the specific bacterial species. Future research should therefore aim to identify and validate the analogous ILA-producing bacteria in the human gut. Certain human-derived *Lactobacillus* strains are also capable of ILA production, including *Limosilactobacillus reuteri* and *Lactobacillus acidophilus* ^31,36^. Other promising candidates include members of the *Bifidobacterium* genus, particularly like *B. infantis* and *B. breve*, which are known to be potent ILA producers and are highly abundant in the human gut, especially in early life ^37^. Second, the upstream mechanism by which *T. spiralis* enriches *L. murinus* remains unresolved. This could involve parasite-induced changes in mucus composition, epithelial nutrients, antimicrobial peptides, bile acids, oxygen tension or type 2 immune remodeling of the intestinal niche. Third, although our data support an AhR-dependent mechanism, the critical responding cell types remain unresolved. Given the broad expression of AhR across epithelial, myeloid, and lymphoid compartments, future studies using cell type-specific AhR deletion will be important for defining where ILA acts most decisively in this model.

In conclusion, our findings define a helminth-remodeled microbial tryptophan metabolic pathway that promotes AhR-dependent immune tolerance and limits inflammatory tissue injury. Rather than acting solely through direct parasite-host interactions, the parasite appears to exploit a microbial metabolic circuit that reinforces host immune restraint and limits excessive inflammatory injury. These findings provide a mechanistic framework for understanding how helminth exposure can shape systemic immune homeostasis and identify the microbial ILA-AhR pathway as a candidate target for postbiotic or microbiota-based strategies aimed at inflammatory disease.

## Acknowledgements

We thank Dr. Dan R. Littman for inspiring our thinking on host–microbiota.

## Author contributions

Conceptualization: X.J., X.L., M.L. and R.S.

Methodology: R.S., Y.L. and Y.W.

Investigation: R.S., N.X. and X.M.,

Visualization: R.S., X.Z. Q.L. and Z.W.

Funding acquisition: X.J., X.L. and M.L.

Writing—original draft: X.J. and R.S.

Writing—review and editing: X.J., X.L., M.L. I.V., P.B. and R.S.

## Materials and methods

### Ethics statement

All animal procedures were approved by the Institutional Animal Care and Use Committee of Jilin University and were performed in accordance with institutional guidelines and the ARRIVE recommendations. C57BL/6J wild-type mice (6-8 weeks old) were purchased from Changchun Yisi Biotechnology Co., Ltd. C57BL/6J AhR-knockout mice (6 weeks old) were obtained from GemPharmatech. Mice were maintained under specific pathogen-free conditions on a 12-hour light/dark cycle with free access to chow and water.

### Helminth infection

*T. spiralis* (ISS534) muscle larvae (ML) were isolated from Wistar rats 35 days after oral infection with 5,000 infective larvae. To generate a helminth infection model, six-week-old male C57BL/6J wild-type (WT) and AhR KO mice were orally inoculated with 250 TsML. Data collection occurred following euthanasia via CO2 exposure at 14 days post-infection (dpi). For the AhR antagonist pretreatment experiments, mice were intraperitoneally injected with 10 mg/kg of CH223191 (Med Chem Express) for 14 days while being infected with *T. spiralis*.

### Antibiotic-treated *T. spiralis* infection

For microbiota-depletion experiments, mice were randomized into three groups (n = 6 per group): antibiotic only, *T. spiralis* only, and antibiotic plus *T. spiralis*. The antibiotic cocktail contained ampicillin (0.5 g/L), neomycin (0.5 g/L), metronidazole (0.5 g/L), and vancomycin (0.25 g/L) and was administered by daily oral gavage from day −14 to 14 dpi. Infected mice received 250 *T. spiralis* muscle larvae by oral gavage. All mice were euthanized at 14 dpi.

### Fecal microbiota transplantation

To test whether helminth-remodeled microbiota were sufficient to transfer the metabolic phenotype, mice were assigned to control, antibiotic-treated, antibiotic plus control-FMT, or antibiotic plus Ts-FMT groups (n = 6 per group). After 28 days of antibiotic treatment, recipient mice received fecal microbiota prepared from uninfected donors or from *T. spiralis*-infected donors at 14 dpi. Fresh fecal pellets were suspended immediately in sterile PBS (1 pellet/mL), homogenized, and centrifuged at 100 x g for 5 minutes at 4℃. The supernatant was used as the transplant inoculum.

### *L. murinus* and ILA administration experiments

To confirm the role of dominant probiotics and key metabolite in helminth infection, *L. murinus* or ILA administration were performed. *L. murinus* (BNCC) was cultured in MRS medium and maintained in an anaerobic incubator using the GasPak 100 system (BD Bioscience) at 37℃. After 21 days of ABX treatment, WT mice and AHR KO mice were gavaged with 200ul of 1×10^9^ CFU/mL *L. murinus* every day for 7 days, and control mice were gavaged with 200 μl of blank MRS medium every day for 7 days. For the ILA treatment experiment, mice were gavaged with either 0.1 mL/20 g of corn oil (Solarbio) or 20 mg/kg of ILA (Med Chem Express) once a day for 7 days after 21 days of ABX pretreatment. For the AhR antagonist pretreatment experiments mice were pretreated with CH223191 at a dosage of 10 mg/kg 1 h before gavage of *L. murinus* or ILA.

### Evaluation of worm burden

To evaluate the effects of *L. murinus* and ILA on *T. spiralis* infection, mice were orally infected with 500 ML. For the *L. murinus* supplementation experiment, mice were gavaged daily with L. murinus (200 μL, 1 × 10^9 CFU/mL) from 1 dpi to 14 dpi, while control mice received 200 μL of sterile blank MRS medium daily during the same period. For the ILA treatment experiment, mice were gavaged daily from 1 dpi to 14 dpi with either corn oil (0.1 mL/20 g body weight) or ILA (20 mg/kg) following oral infection with 500 ML. Intestinal adult worms were collected at 14 dpi.

### LPS -induced cytokine storm model

Acute sepsis models were induced by LPS from Escherichia coli 0111: B4 (L2630, Sigma-Aldrich), LPS dissolved in 200 μl sterile phosphate buffer, and intraperitoneally injected at a dose of 8 mg/kg into each mouse. Other groups of mice were intraperitoneally injected with equal volumes of sterile PBS. For the Ts pretreatment group, each mouse was orally administered 250 TsML and injected with LPS at 14 dpi. For the *L. murinus* and ILA pretreatment groups, each mouse was gavaged daily for 7 days with either 200 μL of *L. murinus* at 1×10⁹ CFU/mL or 20 mg/kg of ILA, followed by LPS injection after the final treatment. For the AhR antagonist pretreatment experiments mice were pretreated with CH223191 at a dosage of 10 mg/kg 1 h before gavage of Ts, *L. murinus* or ILA. Six hours after LPS injection, all mice were euthanized under anesthesia and samples were collected.

### Gene expression analysis

Duodenum and organoid RNA was extracted with TRIzol followed by sequential chloroform, isopropanol, and 75% ethanol treatments under RNase-free conditions. Prime Script RT Master Mix (Takara) served for cDNA synthesis. Quantitative RT-PCR was performed using SYBR Green qPCR Master Mix Kit (Genstar) on a Step One Plus instrument (Applied Biosystems). Gene expression was normalized to β-actin and calculated using the 2^−ΔΔCT^ method. Oligonucleotide sequences are provided in Table S2.

### Lung histology

Lung tissues were immediately fixed in 4% paraformaldehyde (≥24 h), paraffin-embedded, and sectioned at 5 μm. Sections underwent hematoxylin and eosin (H&E) staining prior to light microscopic examination. Lung injury severity was scored per established criteria^38^, assessing four parameters: pulmonary hemorrhage, alveolar congestion, and neutrophil infiltration in vessel walls or airspaces. Lung injury severity was scored 0-4: 0 (minimal), 1 (mild), 2 (moderate), 3 (severe), 4 (maximal). A composite lung injury score was generated by aggregating these four metrics.

### Cytokines assays in serum

Blood and organoid culture supernatants were collected from mice retro-orbital plexuses underwent centrifugation (3000 rpm, 15 min, 4°C), with serum analyzed for cytokines using ELISA kits (Elabscience).

### Flow cytometry

Spleen and mesenteric lymph nodes (MLNs) were aseptically harvested. Single-cell suspensions were generated by gentle mechanical dissociation through cell strainer (70 μm) in ice-cold PBS, followed by cell counting and density adjustment to 1×10⁶/mL. Erythrocyte lysis was performed with red blood cell lysis buffer (Solarbio) for 10 minutes. Cells underwent 4h stimulation at 37℃ in 5% (fetal bovine serum) FBS-RPMI-1640 with PMA/ionomycin cocktail plus protein transport inhibitors (BFA, Biolegend) for Th17 intracellular cytokine staining. Cells were then washed twice in FACS buffer (PBS with 2% FBS), blocked for 15 minutes in FACS buffer containing anti-mouse CD16/32 antibody, and incubated with Fixable Viability Dye eFluor™ 780 and surface antibodies (FITC-anti-CD4, Biolegend; and APC-anti-CD25 antibody, BD Biosciences) at 4℃ for 30 minutes in the dark. Cells were then washed twice in FACS buffer and incubated in Fix/Perm solution (eBioscience) for 30 minutes at 4℃ in the dark. Cells were then washed twice in permeabilization buffer and incubated with intracellular antibodies (PE-anti-IL-17A, BD Biosciences; and PE-anti-Foxp3, eBioscience) for 30 minutes at 4℃ in the dark. Flow cytometric data acquisition used BD FACSCalibur flow cytometer (BD Biosciences), with subsequent analysis in FlowJo software (Tree Star Inc, Ashland, OR).

### Western blot analysis

Proteins were extracted from intestine tissue using PBS buffer containing protease inhibitors. After quantification using BCA Protein Assay Kit, equivalent quantities were resolved on 10-12% gradient SDS-PAGE and electrotransferred to nitrocellulose (NC) membranes. After being blocked in 5% skim milk for 2 h at room temperature, the NC membranes were incubated overnight at 4℃ with primary antibodies (anti-AhR/Cyp1a1 (1:1000) and anti-GAPDH (1:10,000)). Wash the NC membrane with TBST three times, and then incubate it with the secondary antibody at room temperature for 1 h. Finally, the membranes were identified using an enhanced chemiluminescence (ECL) kit.

### Intestinal organoid isolation and treatments

The mice intestinal organoid isolation and treatments was performed as described below. Briefly, Intestinal segments of 5-10 cm were aseptically harvested from 4-6-week-old C57BL/6 wild-type mice. Tissues were longitudinally incised, repeatedly flushed with cold PBS, and villi mechanically scraped using sterile glass slides. Cut the intestine into pieces of 1-2 cm and wash repeatedly with cold PBS. Tissue fragments underwent 30 min incubation in 5 mM EDTA-PBS solution at 4℃ on a rotary shaker. Crypts were subsequently dislodged from the basement membrane by vigorous shaking. Following EDTA dissociation, crypts were enriched in supernatants, filtered through 70 μm cell strainers, and pelleted by centrifugation for 5 min. Intestinal organoids were cultured at 37℃ in a 5% CO2 atmosphere.

For *L. murinus* and ILA treatment experiments, organoids were cultured in *L. murinus* (1×10^6^ CFU/mL), ILA (2 mM), CH223191 (10 μM), LPS (200 μg/mL) for 48 h. The organoids were randomized into five groups before being subjected to LPS experiments: (1) Con. (2) LPS. (3) *L. murinus*+LPS. (4) *L. murinus*+LPS+CH223191 (5). ILA+LPS (6) ILA+LPS+CH223191. For Immunofluorescence experiment, organoids were fixed in 4% paraformaldehyde for 1h at 4℃, dehydrated through 15-30% sucrose for 48 h. Organoids sections were stained with primary antibodies (anti-AhR/Cyp1a1 (1:1000)).

### Molecular Dynamics (MD) Simulation

The crystal structure of the AhR ligand-binding domain was retrieved from the PDB database. Protein and small molecule structure files were imported into GROMACS 2022.3^39^. Topology and simulation boxes were generated using pdb2gmx and gmx editconf, respectively. Energy minimization was performed with the steepest descent algorithm. MD simulations were conducted at 300 K and 1 bar using the Amber99sb-ildn force field. Equilibration consisted of 100,000 steps under the NVT ensemble, followed by 100,000 steps under the NPT ensemble, both with a coupling constant of 0.1 ps and a duration of 100 ps. Production MD simulations were run for 100 ns using gmx grompp and gmx mdrun, during which conformational changes were recorded. Trajectories were analyzed using built-in GROMACS tools to calculate root mean square deviation (RMSD), root mean square fluctuation (RMSF) per residue, and the free energy landscape.

### Surface Plasmon Resonance (SPR) Analysis

The binding affinity between ILA and recombinant human AhR was assessed using a Biacore X100 system with a CM5 sensor chip (Cytiva). AhR protein, diluted in acetate buffer (pH 4.0), was immobilized on the chip following surface activation with EDC and NHS. Serial concentrations of ILA (3.125, 6.25, 12.5, 25, and 50 μg/mL) were prepared in running buffer (PBS containing Tween-20) and injected for kinetic measurements. Sensorgrams were processed and analyzed using the Biacore Evaluation software.

### ILA protection in hACE2 transgenic mice

All K18-hACE2 mice experiments were conducted in the biosafety protection level III Laboratory (ABSL-3) certified by China National Accreditation Service for conformity assessment (CNAS). The experimental work on SARS-CoV-2 virus has been approved by the National Health Commission. K18-hACE2 mice (5-6 weeks old) were randomly assigned to three groups: control group, SARS-CoV-2-infected group, and SARS-CoV-2 plus indole-3-lactic acid group (SARS-CoV-2+ILA). Mice in the SARS-CoV-2 group were intranasally infected with 5.0×10^4^ PFU of SARS-CoV-2 WIV04 strain (IVCAS 6.7512). Mice in the SARS-CoV-2+ILA group received ILA at 20 mg/kg by oral gavage once daily from 7 days before infection to 7 days after infection. Mice in the control and SARS-CoV-2 groups received an equal volume of vehicle. For longitudinal assessment, a subset of mice (n=12) from each group was monitored for 14 days after infection, during which survival and body weight were recorded daily. Body weight change was expressed as the percentage of the initial body weight measured on the day of infection. A separate subset of mice (n=3-5) was euthanized at 6 days post-infection (dpi) for virological, histopathological, and immunological analyses.

Viral titers in lung tissues were determined by plaque assay on Vero E6 cells as previously described ^40^. Briefly, the samples were serially ten-fold diluted using Dulbecco’s modified Eagle’s medium (DMEM) containing 2.5% fetal bovine serum (FBS) plus 2% penicillin/streptomycin, and 100 μL of each dilution was added to Vero E6 cells in 24-well-plates. After 1 h incubation of the plates, the inoculums were replaced by fresh methylcellulose overlay containing 2% FBS. The plates were incubated at 37℃ for 3 d, followed by fixation with 8% paraformaldehyde. Then, the plates were stained with 1% crystal violet, and visible plaques were counted to determine viral titers as plaque-forming units (PFU). For histopathology, mice were euthanized before harvest and fixation of tissues. Lungs were inflated with about 2 ml of 10% neutral buffered formalin using a 3-ml syringe and catheter inserted into the trachea and kept in fixative for 7 days. Tissues were embedded in paraffin, and sections were stained with haematoxylin and eosin. In parallel, lung homogenates were analyzed by ELISA to quantify inflammatory and regulatory cytokines, including TNF-α, IFN-γ, IL-10, TGF-β, and IL-17A, according to the manufacturers’ instructions.

### Metagenomic sequencing and analysis

As for metagenomic sequencing data (PRJNA1321505), raw sequencing reads were processed to obtain valid reads as previously described. Total genomic DNA was extracted from fecal samples using QIAamp DNA Stool Mini Kit (Qiagen). DNA integrity, sizes and concentrations were determined by agarose gel electrophoresis and NanoDrop spectrophotometry (NanoDrop). Sequencing libraries were constructed as previously described. After library quality control, high-throughput sequencing was performed using the NovaSeq6000 platform (Illumina).

Fastp is used for preprocessing raw data from the Illumina sequencing platform to obtain clean data for subsequent analysis^41^. MEGAHIT software is used for assembly analysis of clean data^42^.

With the default parameters, MetaGeneMark is used to perform ORF prediction for scaftigs (>= 500 bp) of each sample^43^, and the information with a length less than 100 nt in the prediction results is filtered out^44^. For the ORF prediction results, CD-HIT software is used to eliminate redundancy^45^ and obtain the nonredundant initial gene catalogue. Clean data of each sample is aligned to the initial gene catalogue by using Bowtie2 to calculate the number of reads of the genes on each sample alignment^46^. Based on the abundance of each gene in the gene catalogue in each sample, basic information statistics, core-pan gene analysis, correlation analysis between samples, and Venn diagram analysis of gene number are performed.

DIAMOND software is used for alignment of unigenes sequences with Micro NR database^47^, which includes sequences from bacteria, fungi, archaea, and viruses extracted from NCBI’s NR database. The alignment is performed useing the blastp algorithm with a parameter setting of 1e^−5 48^. Since each sequence may have multiple alignment results, LCA algorithm applied to systematic taxonomy of MEGAN software was adopted to determine the species annotation information of the sequence^49^. Out of the results of LCA annotation and gene abundance table, the abundance of each sample at each taxonomy and the corresponding gene abundance tables are acquired. The abundance of a species in a sample is equal to the sum of the abundance of those genes annotated as that species. The number of genes of a species in a sample is equal to the number of genes whose abundance is non-zero among the genes annotated as that species. LEfSe software is used for LEfSe analysis.

### Target quantification of tryptophan metabolites

Targeted metabolomic analysis of tryptophan metabolites was performed by Novogene. Co., Ltd. using targeted metabolomics. All standards, including stable isotope-labeled analogs, were sourced from ZZStandards Co., LTD. Methanol (Optima LC-MS), acetonitrile (Optima LC-MS), formic acid (Optima LC-MS) and Ammonium acetate (Optima LC-MS) were purchased from Thermo-Fisher Scientific. Imino-bis (methylphosphonic acid) was purchased from Sigma-Aldrich. Ultrapure water was purchased from Millipore. In brief, the stock solution of individual CCM related compounds was mixed and prepared in CCM related compounds-free matrix to obtain a series of calibrators. The samples were resuspended with liquid nitrogen, and then added to water by well vortexing as the diluted sample. Then 100 μL of them were taken respectively and homogenized with 300 μL of methanol (80%) which contained mixed internal standards by well vortexing. Next, stewing for 10 min. After that, centrifuged at 15,000 g for 15 min. Finally, the supernatant was injected into the LC-MS/MS system for analysis.

The metabolite abundance data were normalized (log-transformation and Pareto scaling) before PCoA to reduce bias from highly abundant metabolites. Heatmaps were generated using the pheatmap package in R (v4.2.1) to visualize differential metabolite patterns across samples. Hierarchical clustering was applied using the Bray-curtis distance metric and complete linkage method.

### Statistical analysis

GraphPad Prism 8.0 was used for statistical analyses and data visualization. Two-group comparisons were performed with unpaired Student’s t tests, and comparisons among three or more groups were performed with one-way analysis of variance as appropriate. For metabolomic and metagenomic analyses, multiple-testing correction was performed with the Benjamini-Hochberg method. A two-sided P value < .05 was considered statistically significant.

### Materials availability

All reagents and resources used in this study should be directed to the lead contact. All reagents will be available based on request and completing a materials transfer agreement.

### Data and code availability

Metagenomic sequencing data are available through the NCBI BioProject database under accession number PRJNA1321505. This study also re-analyzed publicly available COVID-19 metabolomics data from Su et al^29^. No custom code was generated. Additional information required to reanalyze the data reported here is available from the lead contact on reasonable request.

