## Supplementary Information for "Helminth-remodeled microbial indole-3-lactic acid drives AhR-dependent disease tolerance"

**SUPPLEMETARY INFORMATION**

**Supplementary Figures S1-S8**

**Supplementary Tables S1, S2**

**Supplementary Figures**

**Supplementary Fig. S1.**

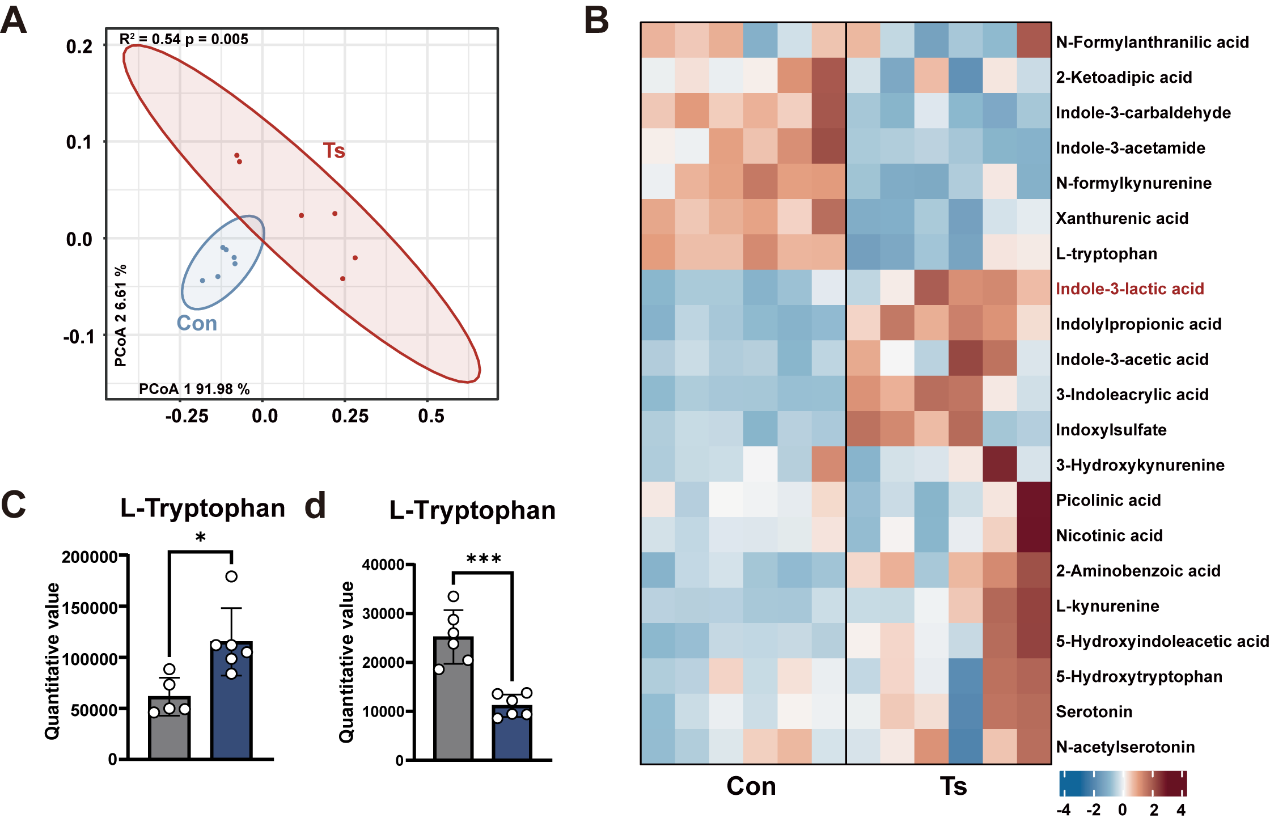

**Figure S1. Ts** **infection remodels host tryptophan metabolism.**

**(A)** Principal Coordinates Analysis (PCoA) showing distinct serum metabolic profiles between control and Ts-infected mice.

**(B)** Heatmap of differentially abundant serum metabolites.

**(C)** The levels of L-Tryptophan in the feces from control and Ts-infected mice.

**(D)** The levels of L-Tryptophan in the serum from control and Ts-infected mice.

Data are presented as mean ± SD. Statistical analysis was performed using an unpaired Student's t-test (c and d). *, p < 0.05, *** p < 0.001.

**Supplementary Fig. S2.**

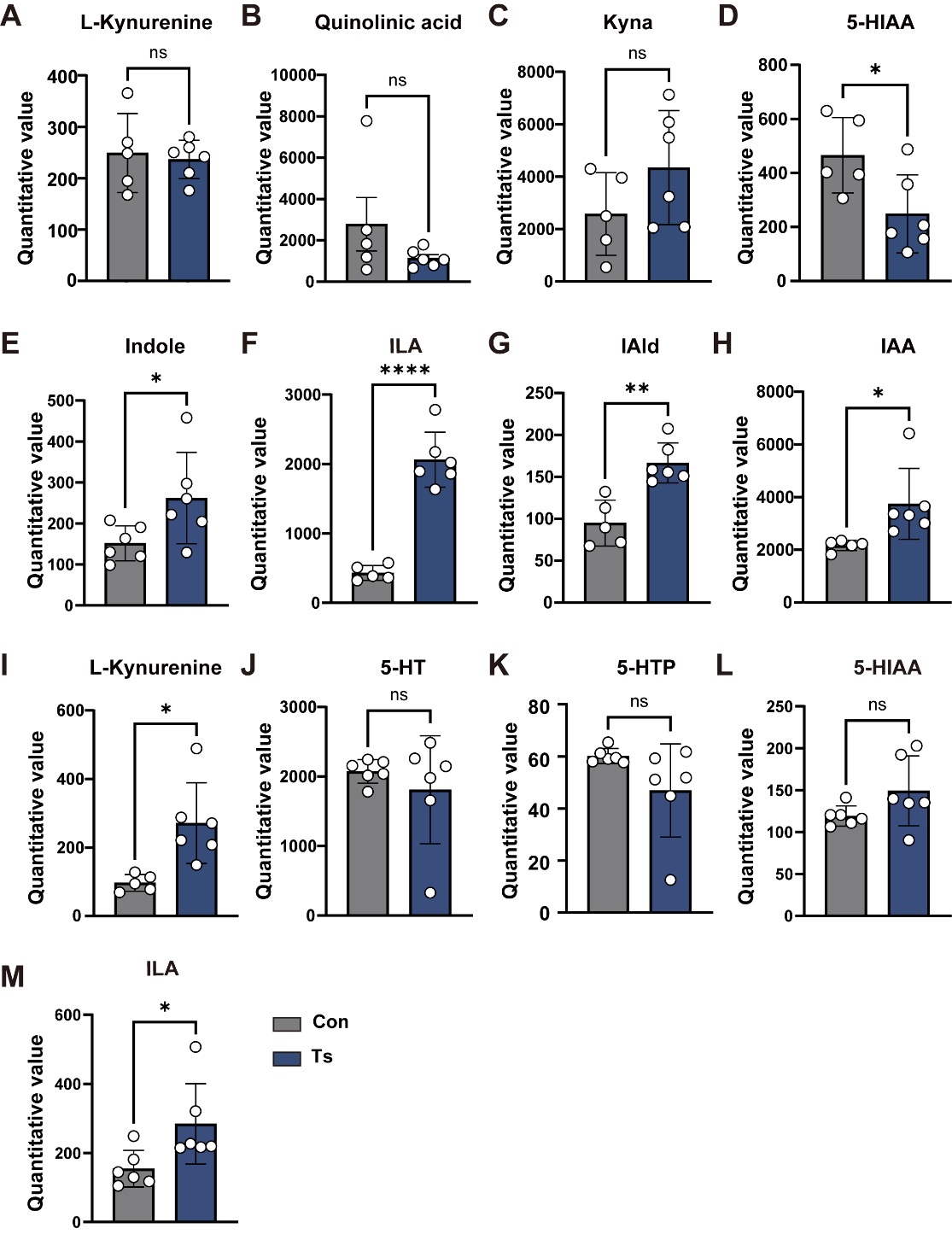

**Figure S2. Ts infection promotes the indole derivative pathway.**

**(A-H)** The levels of tryptophan metabolites in the feces from control and Ts-infected mice.

**(I-M)** The levels of tryptophan metabolites in the serum from control and Ts-infected mice.

Data are presented as mean ± SD. Statistical analysis was performed using an unpaired Student's t-test. ns, no statistical significance, *, p < 0.05, **, p < 0.01, **** p < 0.0001.

**Supplementary Fig. S3.**

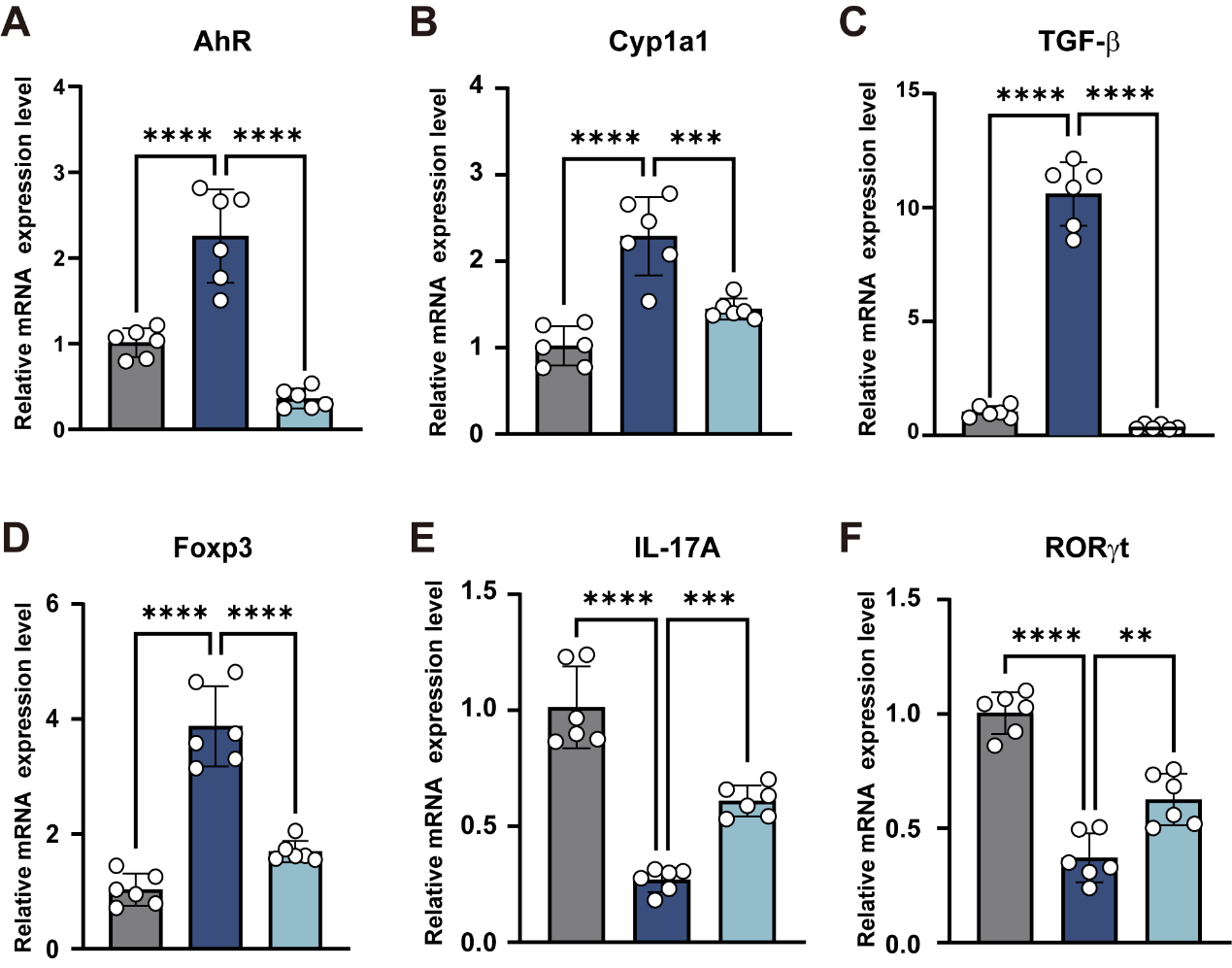

**Figure S3. Ts drives AhR-dependent modulation of the Treg/Th17 balance.**

**(A-F)** qPCR analysis of AhR Cyp1a1 and Treg/Th17-associated genes expression in the intestinal tissue of mice (n=6).

Data are presented as mean ± SD. Statistical significance was determined by one-way ANOVA followed by Tukey multiple comparison test. **, p < 0.01, *** p < 0.001, **** p < 0.0001.

**Supplementary Fig. S4.**

**Fig. 1G, 1I, 1K, 1M**

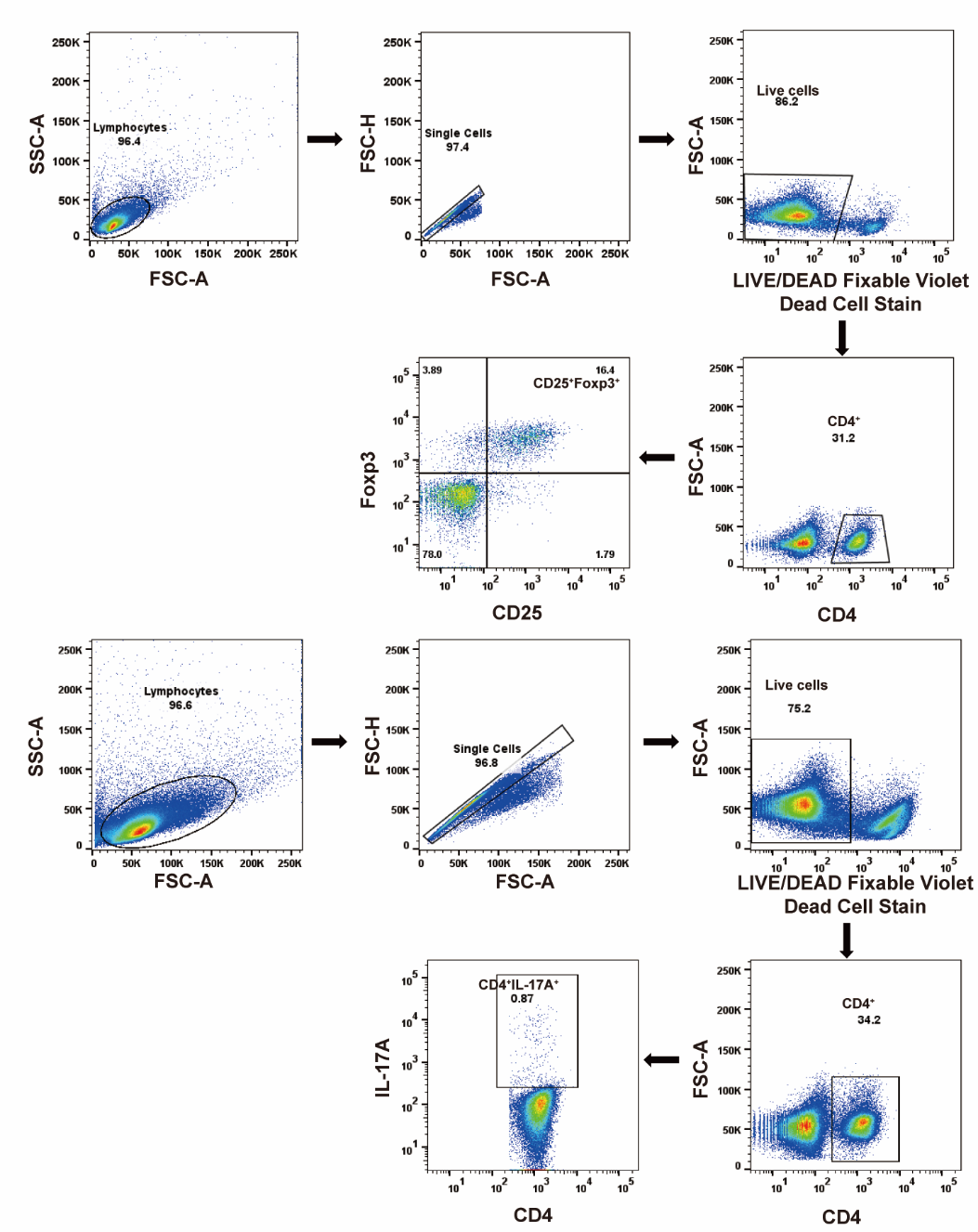

**Figure S4. Flow cytometric gating strategies were used in this study.** Gating strategies for all flow cytometric analyses presented in the main figures of this manuscript.

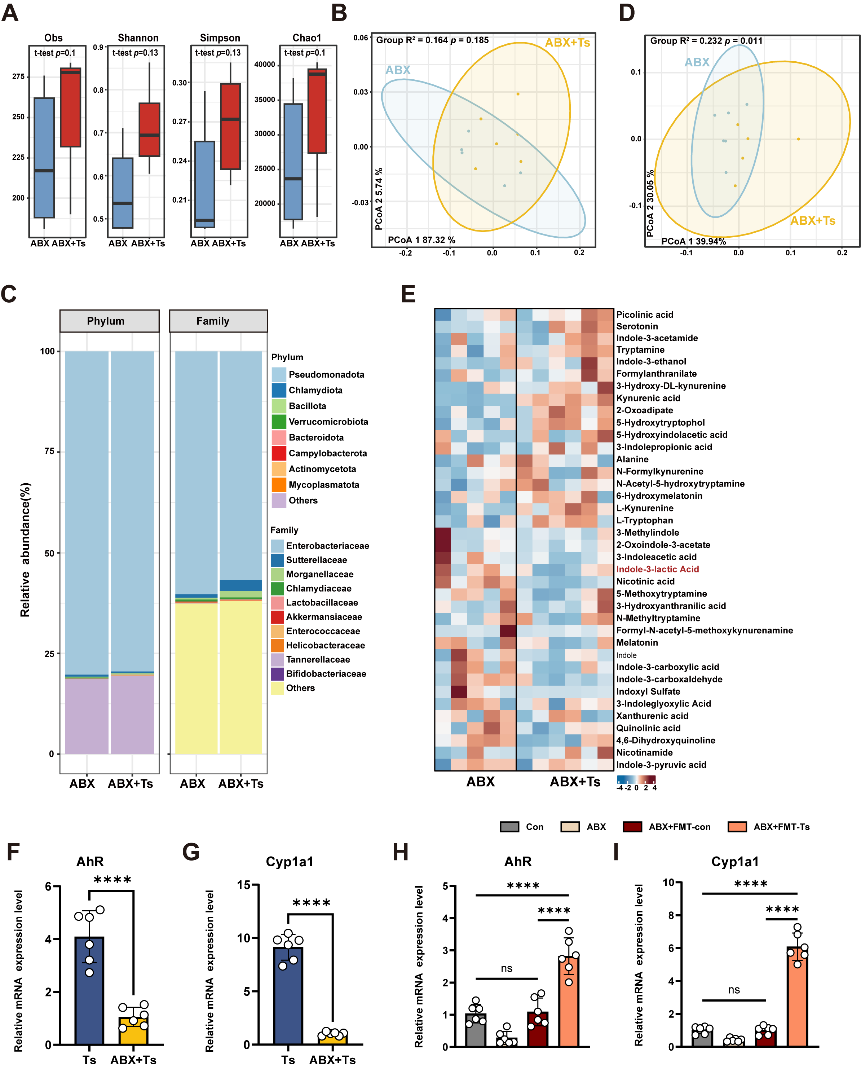
**Supplementary Fig. S5.**

**Figure S5. Ts modulates tryptophan metabolism via the microbiota**

**(A)** α diversity (Obs index, Shannon index, Simpson index and Chao 1) in the microbiota from the ABX and ABX+Ts-infected mice.

**(B)** Principal coordinates analysis (PCoA) plot of metagenomic sequencing from ABX and ABX+Ts-infected mice.

**(C)** Phylum and Family-level median relative abundances.

**(D)** Principal coordinates analysis (PCoA) plot of metabolomic target analyses from ABX and ABX+Ts-infected mice.

**(E)** Heatmap of differentially abundant fecal metabolites.

**(F-I)** qPCR analysis of AhR and Cyp1a1 mRNA expression in the intestinal tissue of mice from different treatment groups. (n=6).

Data are presented as mean ± SD and were analyzed by Student's t-test (F and G) or one-way ANOVA with Tukey's test (H and I). ns, no statistical significance, ****, p <0.0001.

**Supplementary Fig. S6.**

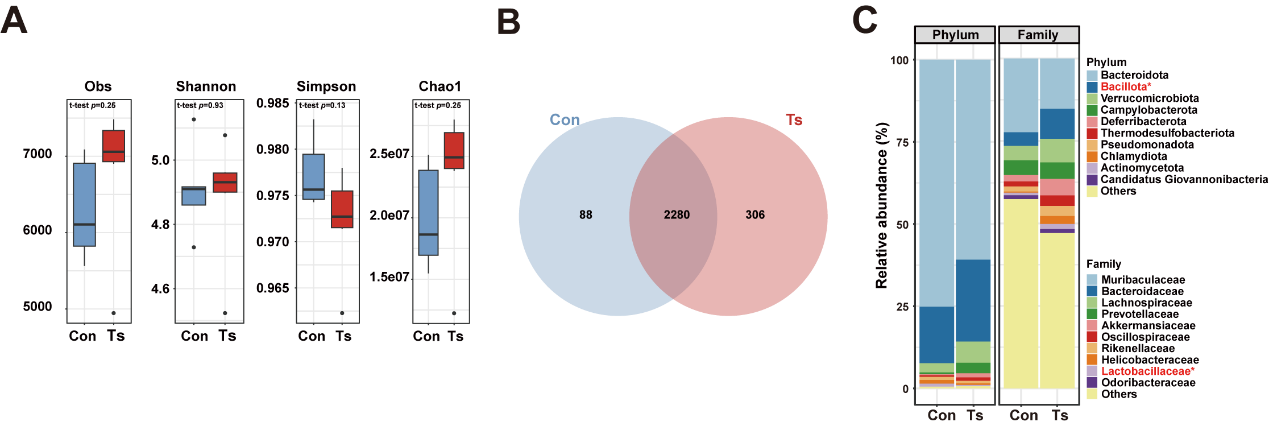

**Figure S6. Ts infection reshapes the host gut microbiota.**

**(A)** α diversity (Obs index, Shannon index, Simpson index and Chao 1) in the microbiota from the controls and Ts-infected mice.

**(B)** Venn of microbiota.

**(C)** Phylum and Family-level median relative abundances.

**Supplementary Fig. S7.**

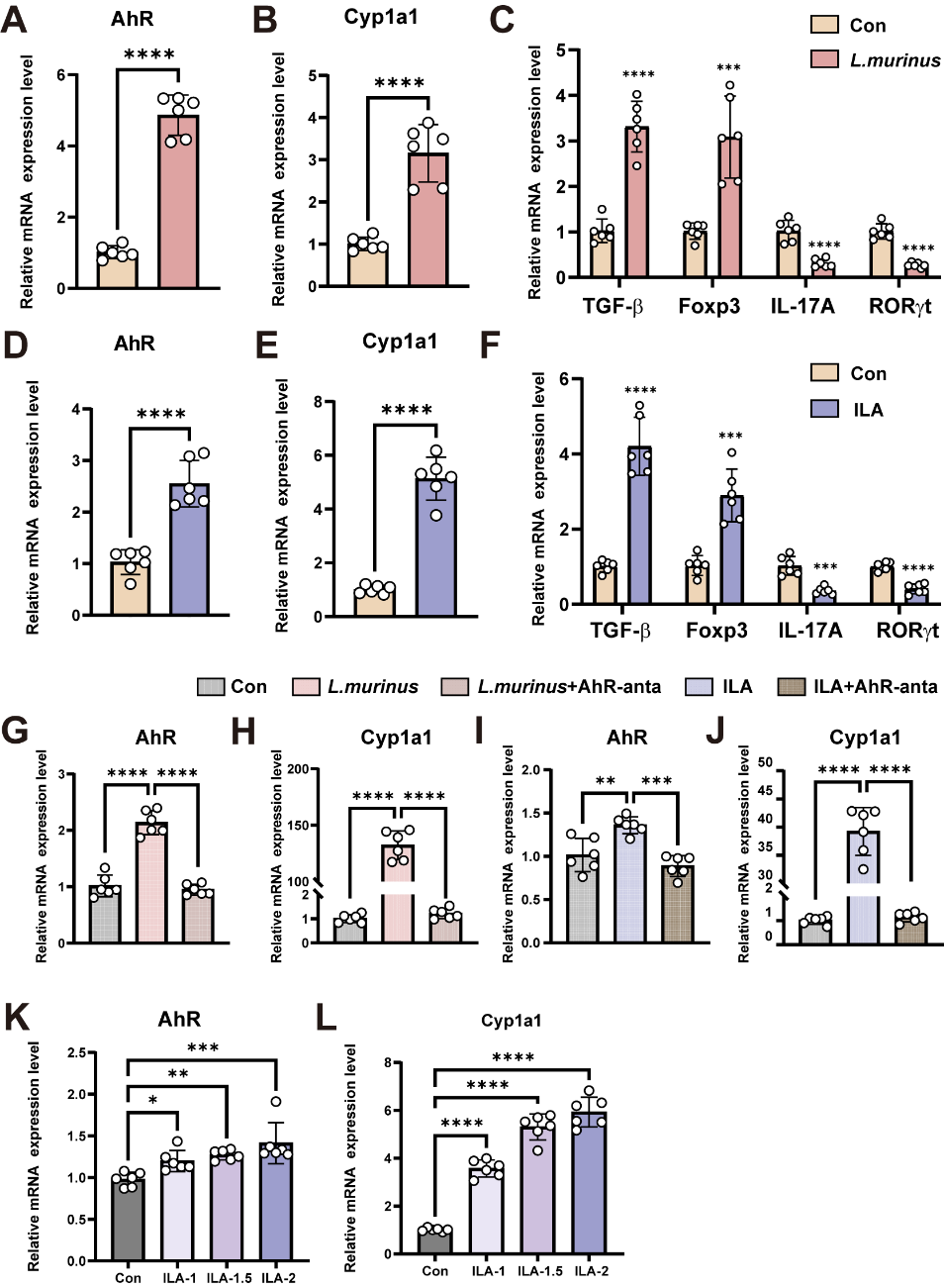

**Figure S7. *L. murinus* and ILA supplementation activates downstream tryptophan metabolism.**

**(A-F)** qPCR analysis of AhR, Cyp1a1 and Treg/Th17-associated genes expression in the intestinal tissue of mice from control, *L.murinus* (A-C) and ILA (D-F) groups (n=6).

**(G-J)** In vitro effects in colonic organoids. Dose-dependent upregulation of AhR and Cyp1a1 mRNA in organoids treated with *L. murinus* (1×10^6^ CFU/mL) or ILA (2 mM) for 48 h, which was markedly attenuated by co-treatment with the AhR antagonist CH223191(10 μM). (n=6)

**(K and L)** qPCR analysis of AhR and Cyp1a1 mRNA expression in the organoids treated with different doses of ILA (n=6).

Data are presented as mean ± SD and were analyzed by Student's t-test (A-F) or one-way ANOVA with Tukey's test (G - L). *, p < 0.05, **, p < 0.01. *** p < 0.001, **** p < 0.0001.

**Supplementary Fig. S8.**

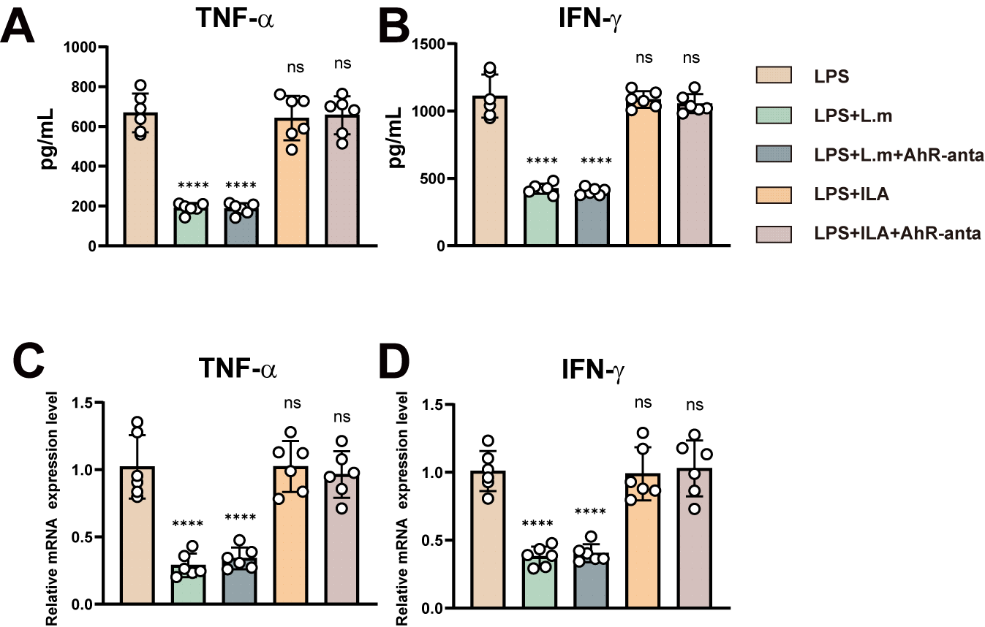

**Figure S8. *L. murinus* and ILA reduce the inflammation of organoids.**

**(A and B)** ELISA analysis of TNF-α, IFN-γ in the organoids culture supernatant (n=6).

**(C and D)** qPCR analysis of TNF-α, IFN-γ mRNA expression in the organoids (n=6).

Data are shown as individual data points and mean ± SD. Statistical significance is calculated using Tukey multiple comparison tests. **** p < 0.0001.

**Supplementary Tables**

**Supplementary Table 1. Key Antibodies, Reagents, and Resources**

| REAGENT or RESOURCE | SOURCE | IDENTIFIER |
| --- | --- | --- |
| Antibodies | | |
| Anti-AhR | Affinity Biosciences | Cat# AF6278; RRID: AB_2839420 |
| Anti-Cyp1a1 | Affinity Biosciences | Cat# DF6882; RRID: AB_2838841 |
| Anti-GAPDH | Affinity Biosciences | Cat# AF7021; RRID: AB_2839421 |
| Anti-AhR | Proteintech | Cat# 67785-1-Ig; RRID: AB_2918549 |
| Anti-Cyp1a1 | Proteintech | Cat# 13241-1-AP; RRID: AB_2877928 |
| CD4(mouse), clone GK1.5 | Biolegend | Cat# 100509; RRID: AB_312712 |
| CD25(mouse), clone PC61 | BD Biosciences | Cat# 561048; RRID: AB_398623 |
| Foxp3(mouse), clone FJK-16s | eBioscience | Cat# 12-5773-82; RRID: AB_465936 |
| IL-17A(mouse), clone TC11-18H10 | BD Biosciences | Cat# 559502; RRID: AB_397256 |
| Bacterial and Virus Strains |  |  |
| Lactobacillus murinus | BNCC | BNCC 194688 |
| Chemicals, Peptides, and Recombinant Proteins | | |
| Indole-3-Lactic Acid (ILA) | MedChemExpress | Cat# HY-113099, CAS: 1821-52-9 |
| CH223191 | MedChemExpress | Cat# HY-12684, CAS: 301326-22-7 |
| Ampicillin | Sigma | Cat# A5354, CAS: 69-52-3 |
| Neomycin | Sigma | Cat# N6386, CAS: 1405-10-3 |
| Metronidazole | Sigma | Cat# 16677, CAS: 443-48-1 |
| Vancomycin Sigma | Sigma | Cat# V2002, CAS: 1404-93-9 |
| Man-Rogosa-Sharpe (MRS) broth | BNCC | BNCC 363097 |
| Cell Activation Cocktail | Biolegend | Cat# 423302 |
| Brefeldin A | eBioscience | Cat# 00-4506-51 |
| RPMI 1640 medium | ThermoFisher | Cat# 22400089 |
| Fetal Bovine Serum | ThermoFisher | Cat# 10091148 |
| TRNzol Reagent | Sigma | Cat# 15596018CN |
| 2×RealStar Fast SYBR qPCR Mix (Low ROX) | Genstar | Cat# A304-05 |
| IntestiCultTM organoid growth medium | StemCell | Cat# 06005 |
| Matrigel matrix, growth factor reduced | Corning | Cat#356231 |
| **Continued** |  |  |
| REAGENT or RESOURCE | SOURCE | IDENTIFIER |
| 0.5M EDTA PH8.0 | Beyotime | Cat# C0196 |
| Red Blood Cell Lysis Buffer | Solarbio | Cat# R1010 |
| Recombinant Human AHR | Cloud-clone | Cat# RPB354Hu01 |
| LPS | Sigma | Cat# L2630 CAS: 93572-42-0 |
| Corn oil | Solarbio | Cat# IC9001 CAS: 8001-30-7 |
| CM5 sensor chip | Cytiva | Cat# 29104988 |
| Critical commercial assays | | |
| Mouse TNF-α ELISA assay kit | Elabscience | E-MSEL-M0002 |
| Mouse IFN-γ ELISA assay kit | Elabscience | E-MSEL-M0007 |
| Mouse IL-17A ELISA assay kit | Elabscience | E-MSEL-M0006 |
| Mouse IL-10 ELISA assay kit | Elabscience | E-MSEL-M0031 |
| TGF-β ELISA assay Kit | Elabscience | E-EL-0162 |
| Cytofix/Cytoperm Fixation/Permeabilization Kit | BD Biosciences | Cat# 554714 |
| Foxp3 transcription factor staining kit | eBioscience | Cat# 00-5523-00 |
| PrimeScript FAST RT kit with gDNA Eraser | Takara Bio | Cat# RR092A |
| Deposited data | | |
| Metagenomic sequencing data | This paper | NCBI BioProject: PRJNA1321505 |
| Experimental Models: Organisms/Strains | | |
| Mouse: C57BL/6J | Changchun Yisi Biotechnology Co., Ltd. | N/A |
| Mouse: C57BL/6JGpt-Ahrem1Cd1980/Gpt | GemPharmatech | Strain# T012419 |
| Oligonucleotides | | |
| See Table S2 for primers |  |  |
| Software and algorithms | | |
| Prism v8.0c | GraphPad Software | https://www.graphpad.com/scientificsoftware/prism/ |
| ImageJ | ImageJ | https://imagej.net/software/imagej/ |
| FlowJo v10 | BD Biosciences | https://www.flowjo.com/ |
| R v4.2.1 |  | https://www.r-project.org/ |

**Supplementary Table 2. qRT-PCR primers were used in this study.**

| **Target genes** | **Forward primer sequence (5´→ 3´)** | **Reverse primer sequence (5´→ 3´)** |
| --- | --- | --- |
| AhR | GAGCACAAATCAGAGACTGG | TGGAGGAAGCATAGAAGACC |
| Cyp1a1 | GTTAACCATGACCGGGAACT | GTGACCTTCTCACTCAAGCG |
| TGF-β | TGATACGCCTGAGTGGCTGTCT | CACAAGAGCAGTGAGCGCTGAA |
| Foxp3 | GGCAGAGGACACTCAATGAAAT | TCTCCACTCGCACAAAGCAC |
| IL-17A | TACCTCAACCGTTCCACGTC | TTTCCCAACCGCATTGACACA |
| RORγt | CGAGATGCTGTCAAGTTTGG | CACTTGTTCCTGTTGCTGCT |
| β-actin | GGCTGTATTCCCCTCCATCG | CCAGTTGGTAACAATGCCATGT |
